# Cancer pathway connectivity resolved by drug perturbation and RNA sequencing

**DOI:** 10.64898/2025.12.11.692401

**Authors:** Thi Huong Lan Do, Caroline Lohoff, Ferris Jung, Sandra Kummer, Marcel Fabian Pohly, Sebastian Scheinost, Jennifer Huellein, Vladimir Benes, Wolfgang Huber, Junyan Lu, Thorsten Zenz

## Abstract

Pathway inhibitors are a backbone of cancer treatment. The configuration of pathway dependencies varies from tumour to tumour. Better treatment for individual patients could be designed if the pathway wiring were readily measurable.

Here, we characterise the transcriptional responses of 116 lymphoma patient samples exposed to ten drug perturbations. We used factor analysis to decompose individual and shared drug effects, thereby generating a pathway connectivity map of chronic lymphocytic leukemia (CLL). The expression profiles of the major disease subgroups, defined by IGHV mutation status, became more similar to each other after BTK inhibition, consistent with B-cell receptor (BCR) signalling as a driver of their phenotypic difference. An even stronger convergence was observed with combined IRAK4 and BTK inhibition, indicating cooperation of BCR and toll-like receptor (TLR) signalling in CLL. We identified genetic aberrations (*BRAF*, *TP53*, deletion 17p, deletion 15q, trisomy 12) that modulated drug effects in CLL and constituted specific interaction patterns. IRAK4 inhibition effects depended on the presence of trisomy 12, a finding that suggests that the trisomy 12 driver event in CLL acts by gene dosage-dependent IRAK4 upregulation and amplification of the BCR/TLR cooperation.

Our results highlight the potential of systematic drug perturbation assays with transcriptome readout to map pathway interconnectivity and functionally annotate tumour drivers.

## Introduction

Chronic lymphocytic leukemia (CLL) is a common B-cell lymphoma in adults. More than 200 driver genes and chromosomal aberrations are known, which target a broad range of signalling pathways including the B-cell receptor (BCR), DNA damage response and NOTCH pathway^1–4^. A more precise understanding how this broad range of pathways cooperate and contribute to the CLL phenotype should facilitate better treatments. Multi-omics profiling of CLL patient samples has revealed major biological dimensions of CLL, such as the epigenetic cell-of-origin fingerprint^5,6^, the clinical and biological impact of IGHV mutations^7,8^, autonomous BCR signalling^9^ and CLL proliferative drive (CLL-PD)^8,10–13^. Nonetheless, many common mutations and copy number variants (CNVs), e. g. trisomy 12, have not yet been conclusively linked to a function^10^.

The ability to measure multi-omics data in large cancer cohorts has refined molecular subgrouping, improved outcome prediction and treatments^12,14,15^. Recently, however, progress has often focused on the discovery of rarer disease variants with the inherent downside that findings only apply to a small proportion of patients^1,3^. Furthermore, clinical precision oncology has largely relied on cancer genome sequencing and the identification of genetic mutations as biomarkers^16,17^, while other omics data have remained research tools. Among these, gene expression profiling is particularly suited for the detailed characterisation of drug effects. This concept has been applied in the “Connectivity Map”, which represents a landmark study mapping over 80,000 genetic and chemical perturbations to gene expression signatures in 29 cancer cell lines^18,19^, and has been widely used to identify drug effects, drug repurposing and construct gene regulatory networks^18,20–22^. By using the perturbation signatures, transcriptional consequences can specifically be linked to the targeted upstream cause and therefore facilitate the understanding of cellular mechanisms^18^.

In this study, we applied a set of 10 small-molecule drugs to a collection of 116 primary tumours, to investigate how these drugs work, capture the wiring of targeted pathways and identify modulators (i.e. mutations) of drug response.

## Results

### Shallow depth transcriptional profiles capture CLL disease dimensions and drug effects

In order to increase feasibility and cost-effectiveness of profiling hundreds to thousands of samples, we first compared gene expression from five CLL samples profiled with whole-transcript deep (TruSeq) and 3’-end-based library preparation with shallow depth RNA sequencing (QuantSeq^23^) (Suppl. Fig. 1a-d). Although the sequencing depth was reduced up to 30-fold (2.2-3.8 vs. 43-73 million reads per sample), QuantSeq recovered 71% of the genes detected by TruSeq with reproducible quantification (*R* = 0.82-0.85) (Suppl. Fig. 1e-g).

We selected a cohort of 108 chronic lymphocytic leukemia (CLL) and eight other B-cell Non-Hodgkin (B-NHL) lymphoma patient samples with available molecular profiles and clinical data (Fig. 1) and used QuantSeq to generate 1,086 (1,010 for CLL, 76 for B-NHL; 92% successfully prepared libraries) gene expression profiles after 48h of *ex-vivo* treatment with inhibitors used clinically (BTK inhibitor ibrutinib, PI3K inhibitor duvelisib, MEK inhibitor trametinib, mTOR inhibitor everolimus, XPO1 inhibitor selinexor), compounds targeting key signalling pathways (IRAK4 inhibitor compound 26, AKT inhibitor MK2206, MDM2 inhibitor and p53 activator nutlin-3a, BET inhibitor IBET762), and one combinatorial drug treatment (BTK and IRAK4 inhibition) (Fig. 1a, Suppl. Fig. 2a,b, Suppl. Table 1). IGHV mutational status, trisomy 12 and *TP53* mutation influenced viability of cells after drug treatment consistent with our prior findings^11^ (Suppl. Fig. 2c-e).

**Fig. 1:**
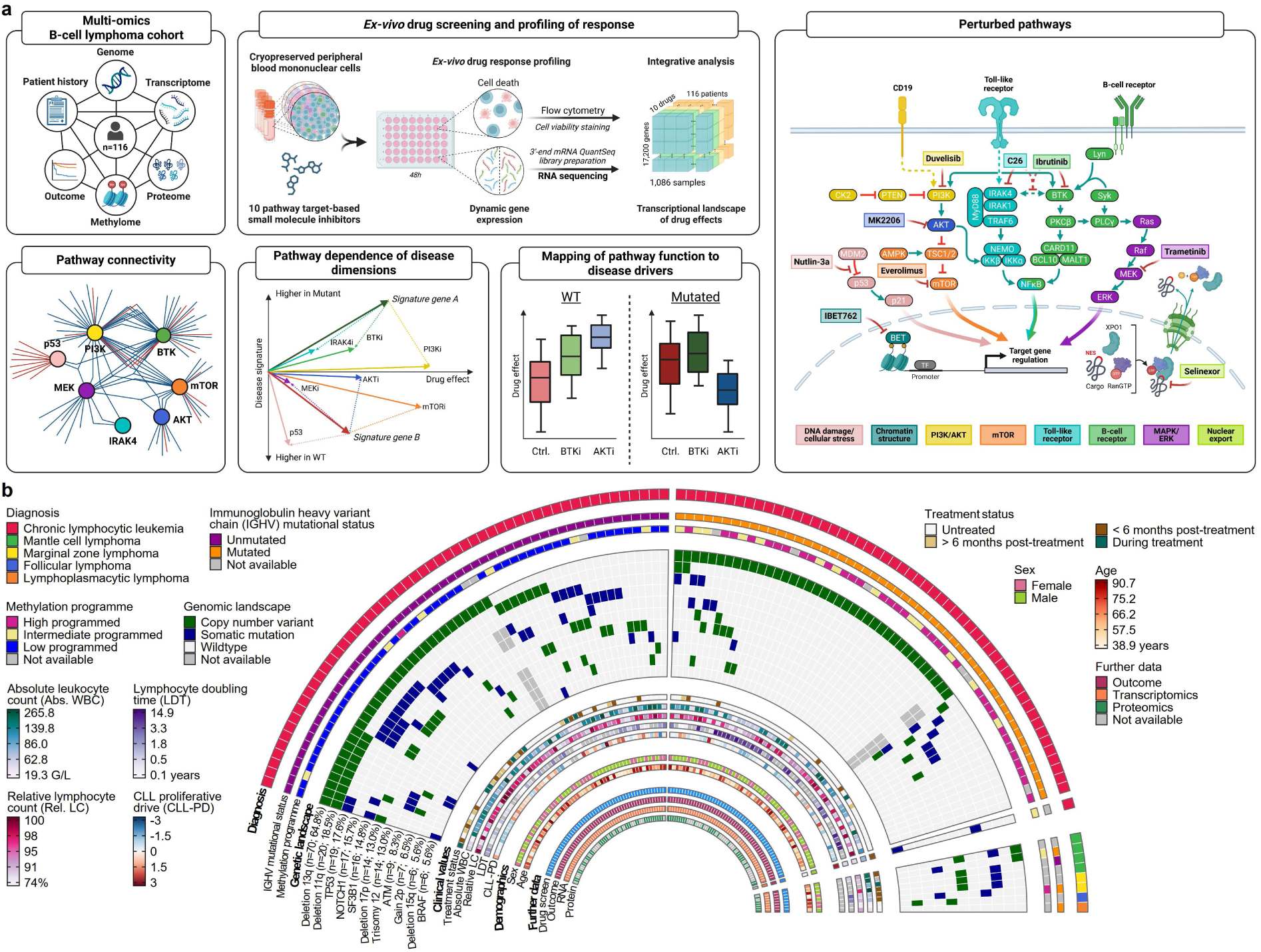
Study overview. **a**, Study outline (https://BioRender.com/6i76n18). **b**, Genetic landscape of study cohort including recurrent genetic aberrations (> 5 cases), clinical and demographic parameters, and availability of additional molecular omics profiling data. Colours for continuous variables are scaled by minimum value, 25th, 50th, 75th percentile and maximum value. Numbers indicated in brackets show case number and frequency of patients with respective aberration for the CLL cohort.

We began our analysis by performing principal component analysis (PCA) to investigate which gene expression programmes, as characterised by factors (≙ principal components), were the main contributors of transcriptional variation in our data set. PCA of the DMSO-treated control samples showed that transcriptional variation between patients was predominantly driven by disease type, IGHV, methylation group^5,6^ and trisomy 12 status (Suppl. Fig. 2f). This result was highly similar to results from studies using conventional RNA sequencing^13^ and shows that we were able to capture the main biological disease dimensions after *ex-vivo* culture and markedly reduced sequencing depth (Suppl. Fig. 2g,h). PCA of all samples including drug treatments revealed that the first component (PC1) captured a contribution of sample viability to the transcriptional variation independent of perturbation (Fig. 2a,b, Suppl. Fig. 2i, Suppl. Fig. 3a-c). This viability signature was enriched in pathways linked to a general upregulation of hypoxia, p53 pathway, TNFα signalling via NFκB, and downregulation of MYC targets and OXPHOS preferentially in drug-perturbed samples (Suppl. Fig. 3d). Variation driven by IGHV status/methylation programme and trisomy 12 were reflected by PC2 and PC3 (Fig. 2c, Suppl. Fig. 2i,j). We also used *t*-Distributed Stochastic Neighbour Embedding (*t*-SNE) to visualise samples’ gene expression similarities and found patterns that were dominated by inter-patient differences and inherent molecular disease features (Fig. 2d-e). After subtracting the patient-specific baseline signatures, the embeddings captured drug specific-effects (Fig. 2f, Suppl. Fig. 2k).

**Fig. 2:**
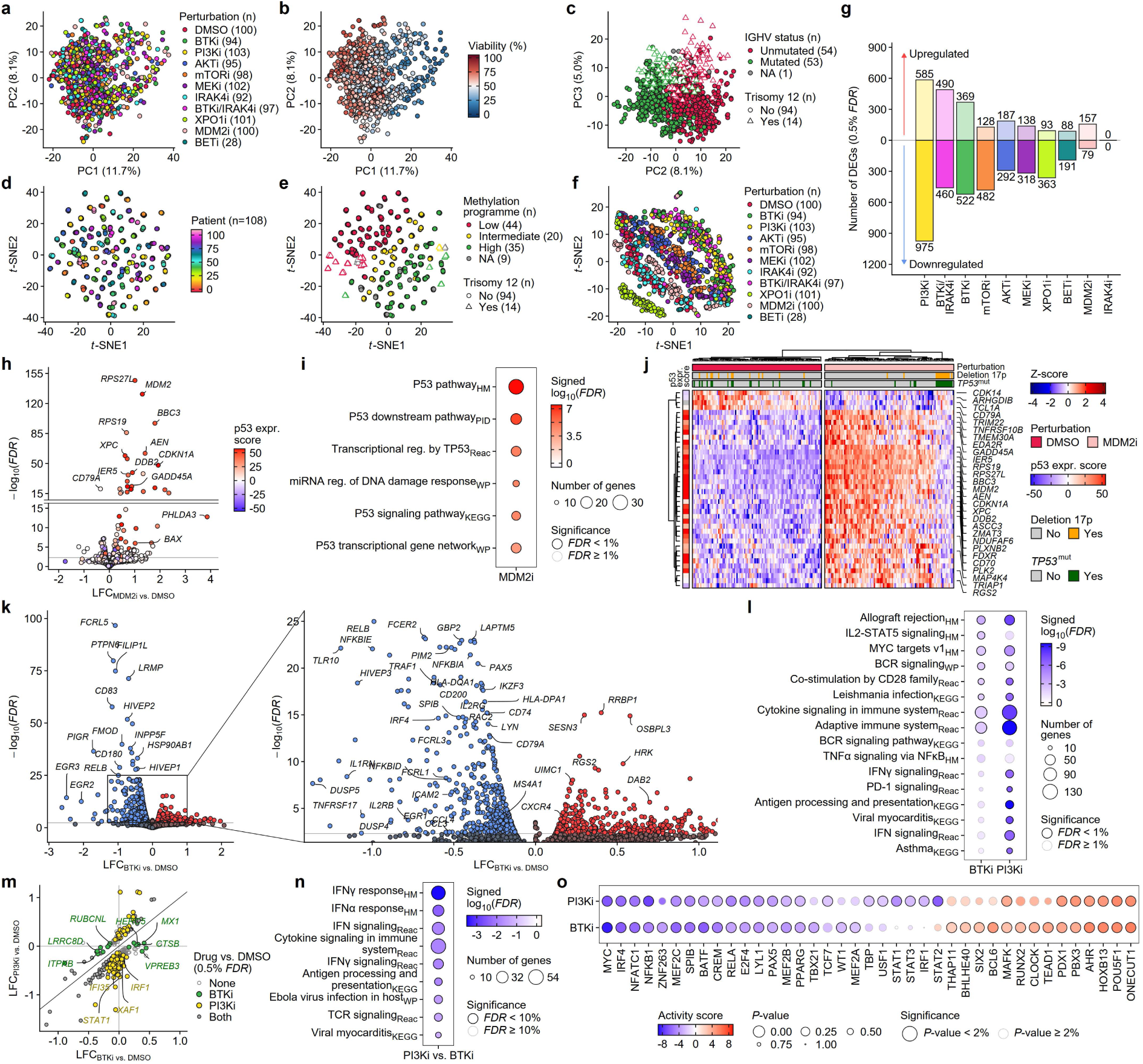
Transcriptional landscape of drug effects in CLL. **a**, First two principal components of PCA based on the 1,000 most variant genes coloured by perturbation and **b**, absolute viability of each drug-treated sample measured by flow cytometry. **c**, PC2 and PC3 of PCA based on the 1,000 most variant genes coloured by IGHV and trisomy 12 status. **d**, *t*-SNE plot based on the 1,000 most variant genes coloured by patient and **e**, by methylation groups. Samples from the same patient form tight clusters. **f**, *t*-SNE plot based on the 1,000 most variant genes after adjustment for the patient effect coloured by perturbation. **g**, Number of differentially expressed genes (DEGs) between drug- and DMSO-treated samples (0.5% *FDR*, |fold change| > 1.05). Induction or suppression for each treatment are shown. **h**, Volcano plot summarising the differential gene expression after MDM2 inhibitor treatment. Genes are annotated according to p53 expression score^50^. **i**, GSEA with DEGs after MDM2 inhibitor treatment (0.5% *FDR*) using hallmark and canonical pathways gene sets. Gene sets were filtered by *FDR* (< 1%) and number of enriched genes (> 10). **j**, Heatmap with hierarchical clustering showing patient effect-adjusted normalised expression of top 40 DEGs after MDM2 inhibitor treatment. *TP53* mutation and deletion status are annotated. **k**, Volcano plot and zoom-in summarising the differential gene expression after BTK inhibitor treatment. **l**, GSEA with DEGs after BTK and PI3K inhibitor treatment (0.5% *FDR*) using hallmark and canonical pathways gene sets. Gene sets were filtered by *FDR* (< 1%) and number of enriched genes (> 10). Depicted are the top 10 gene sets for BTK and PI3K inhibition. **m**, Scatter plot of drug effect sizes for DEGs between PI3K and BTK inhibitor treatment (10% *FDR*). Colours and shapes depict in which comparisons the gene was differentially expressed (0.5% *FDR*). **n**, GSEA with DEGs between PI3K and BTK inhibitor treatment (10% *FDR*) using hallmark and canonical pathways gene sets. Gene sets were filtered by FDR (< 10%) and number of enriched genes (> 10). **o**, Inferred transcription factor (TF) activity using all genes. TFs were filtered by *FDR* (< 10%) and number of targets in TF regulon (> 30 target genes).

### Individual drug effects

To understand the effect of individual drugs, we extracted drug-specific transcriptional signatures by performing differential gene expression analysis between drug- and control-treated samples (Fig. 2g). We first assessed the response to nutlin-3a, a MDM2 inhibitor and activator of the transcription factor p53^24^. We identified the p53 activation signature in primary CLL cells, which consisted of a strong upregulation of well-known transcriptional p53 targets and pathways including *BAX*, *MDM2* and *CDKN1A*/p21 (Fig. 2h-i). This response was blunted in cases with *TP53* mutation and/or deletion in chromosome 17p (Fig. 2j).

Inhibition of PI3K and BTK elicited the strongest responses, as assessed by the number of differentially expressed genes, in line with the role of BCR signalling as a key driver of CLL^7–9^ (Fig. 2g). BTK inhibition led to the downregulation of genes associated with B-cell activation and function (*FCRL5*, *FILIP1L*, *MS4A1*/CD20, *CCL3*/*4*) and phosphatases (*PTPN6*, *DUSP4/5*) (Fig. 2k,l). We also found downregulation of key B-cell transcription factors (*PAX5*, *IRF4*, *IKZF3*) and members of the NFκB pathway (*REL*, *RELB, NFKB1),* suggesting a role of active BCR signalling in the maintenance of the regulatory B-cell identity programme. This signature overlapped with reported transcriptional signatures after BCR stimulation and *in-vivo* BTK inhibitor treatment^25–27^ (Suppl. Fig. 4a).

PI3K inhibition led to the downregulation of gene sets enriched in immune cell response, cytokine signalling, interferon (IFN) response and BCR signalling, similar to the effects upon BTK inhibition but more pronounced (Fig. 2l,m, Suppl. Fig. 4b,c). We investigated the similarities and differences between BTK and PI3K inhibition by directly comparing the effects of the two inhibitors against each other (Fig. 2m, Suppl. Fig. 4d). Genes only modulated by the BTK inhibitor included components of B-cell development and function, endocytosis and autophagy, and the IFN response pathway. PI3K inhibition exclusively downregulated genes associated with IFN and cytokine signalling (Fig. 2n, Suppl. Fig. 4d). To understand whether these differences were driven by a common upstream mediator, we inferred the transcription factor activities from the differentially expressed genes^28^ in which transcription factor activity of STAT family members and in particular gene expression of *STAT1* was downregulated by PI3K but not BTK inhibition (Fig. 2o, Suppl. Fig. 4e-f).

Inhibition of AKT modulated gene sets involved in antigen processing and presentation, IFN signalling and suppression of MHC class II molecules similar to the PI3K inhibitor signature (Suppl. Fig. 5a,b). Differentially expressed genes between PI3K and AKT inhibitor treatment mostly consisted of BTK-dependent targets that were only suppressed upon PI3K inhibition (Suppl. Fig. 5c). The mTOR inhibitor signature showed overlap with the AKT and PI3K inhibitor signatures in the context of antigen processing and adaptive immune system (Suppl. Fig. 5b,d), and specifically included suppression of gene sets associated with lysosome, cholesterol homeostasis and mTORC1 signalling (Suppl. Fig. 5b,g). The predominant mechanisms of action (MoA) induced by the MEK inhibitor were cytokine signalling and IFN response (Suppl. Fig. 5b,e-g).

Bromodomain and extra-terminal motif (BET) proteins are readers of acetylated histone marks and regulate gene expression^29^. Inhibition of BET proteins led to upregulation of genes associated with translation, ribosome and metabolism of RNA (Suppl. Fig. 5h,i). Selinexor is a small-molecule that prevents the translocation of proteins from the nucleus to the cytoplasm by inhibiting the nuclear exporter XPO1^30^. *XPO1* was among the top upregulated genes together with solute carriers, nucleotide- and protein-binding adapters, and members of the zinc finger protein family (Suppl. Fig. 5j,k). Gene expression of p53 was also increased after XPO1 inhibitor treatment without induction of the transcriptional p53 activation signature (Suppl. Fig. 5l). Inhibition of toll-like receptor (TLR) signalling with the IRAK4 inhibitor did not lead to significant differential gene expression (Fig. 2g).

### Connectivity of signalling and perturbation effects in CLL

The use of multiple drugs allowed us to understand how important signalling axes in CLL are connected. For this, we constructed a pathway connectivity network as a bipartite graph in which treatment and gene nodes are connected by edges when the perturbation led to differential expression of that gene. We found that the BTK and PI3K inhibitors were strongly linked through their common BCR-inhibiting MoA. The AKT inhibitor formed close connections to the PI3K and mTOR inhibitors along the PI3K/AKT/mTOR pathway (Fig. 3a). We also illustrated these relationships by comparing the perturbations directly against each other (Fig. 3b). Because drug perturbations typically do not affect a single gene in isolation, but simultaneously modulate the expression of multiple related genes, we used a factorisation technique, namely guided sparse factor analysis^31^ (GSFA) to decompose the full set of drug-induced differentially expressed genes into individual latent factors of co-regulated genes (Fig. 3c-e, Suppl. Fig. 6). We identified drug-specific effects upon inhibition of MDM2 (factor 18), XPO1 (factor 12), BET (factor 16) and MEK (factor 14), but also found modules affected by multiple drugs at variable intensities (Fig. 3c,d, Suppl. Fig. 6a,b). These shared axes of pathway heterogeneity distinguished canonical BCR signalling (factor 8; BTK/PI3K inhibition) from antigen processing (factor 17; PI3K/AKT inhibition), and the lysosome pathway (factor 20; mTOR/MEK inhibition) (Fig. 3c,e, Suppl. Fig. 6b,c). We also identified modules regulated in opposing directions, which could indicate compensatory pathway activation, and link them to IFN (factor 14; MEK/XPO1 vs. mTOR/BTK inhibition) and lysosomal signalling (factor 20; MEK/mTOR vs. PI3K/BTK inhibition) (Fig. 3c-e). Next, we used linear regression to identify the impact of the most prevalent disease aberrations and CNVs on individual patient factor contributions. Mutation or deletion of *TP53* was negatively associated, while trisomy 12 was positively associated with the response to the MDM2 inhibitor captured in factor 18 (Suppl. Fig. 6d). BTK and MEK inhibitor responses reflected in the factors 17 and 14, respectively, were positively associated with unmutated IGHV and trisomy 12 (Suppl. Fig. 6e-f).

**Fig. 3:**
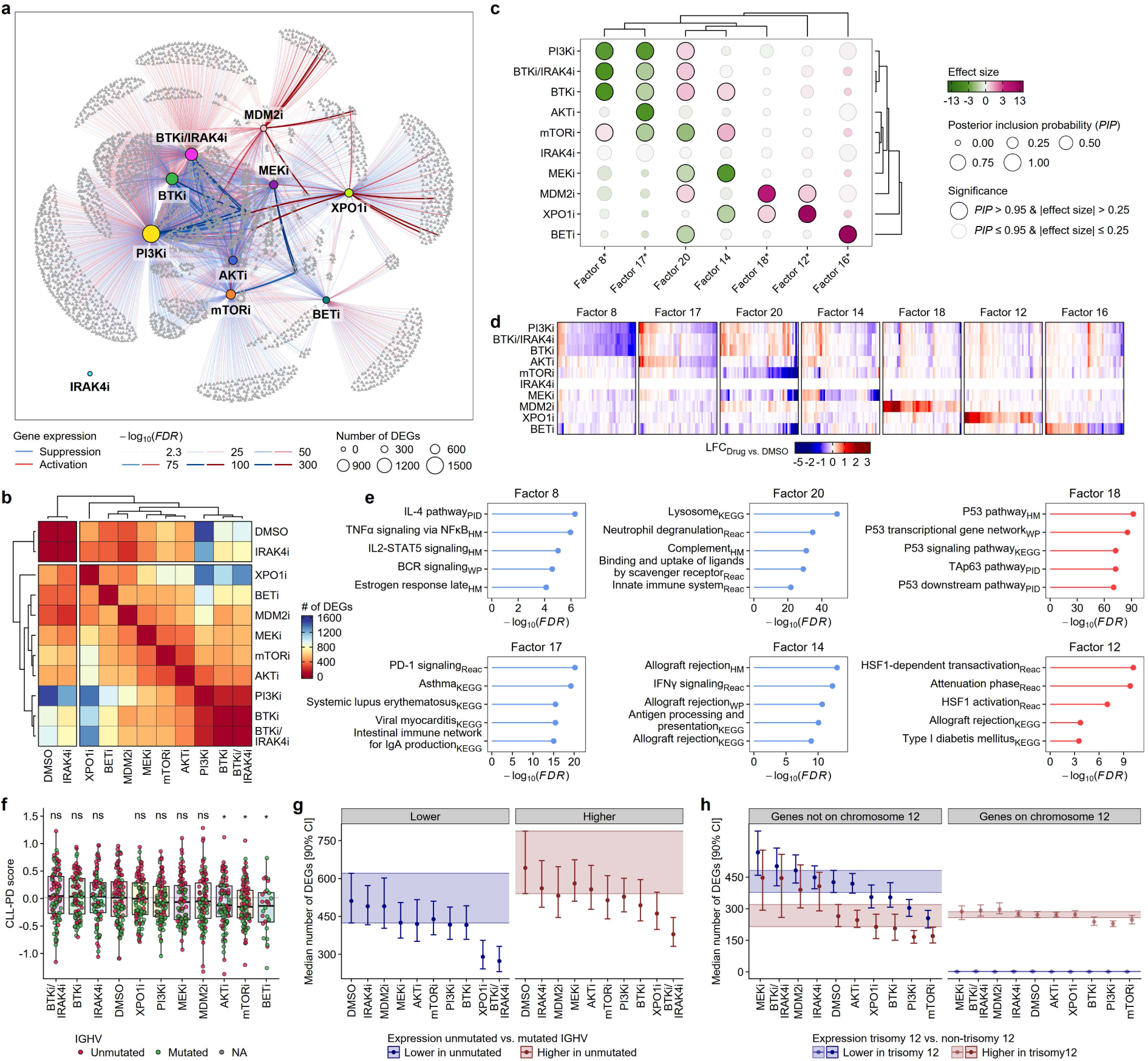
Pathway connectivity in CLL. **a**, Pathway connectivity network shown as a bipartite graph. Genes and perturbations are connected if the perturbation led to differential expression of that gene. The total number of DEGs for each treatment is depicted by the size of the central nodes. Colour and thickness of edges represent whether genes are induced or suppressed, and significance of drug effect, respectively. **b**, Heatmap with hierarchical clustering showing number of DEGs after pairwise drug-drug comparison (0.5% *FDR*). **c**, Estimated effects of drug perturbations on co-regulated gene modules represented by latent factors. The posterior inclusion probability (*PIP*) denotes the probability that a perturbation affects the factor loading. Factors with significant loadings (*PIP* > 0.95, |effect size| > 0.25) are shown. Asterisks denote factors for which the sign was inverted to reflect the direction of the drug effects. **d**, Heatmap showing the drug effects for 50 genes with the highest gene weights in each factor. **e**, GSEA with the gene weights for each factor using hallmark and canonical pathways gene sets. Gene sets were filtered by *FDR* (< 1%) and number of enriched genes (> 10). The sign of weights was reverted in accordance to (c). **f**, CLL-PD scores computed from the transcriptomics data for each individual treatment. Significance levels were calculated using two-tailed paired Student’s *t*-test with DMSO as reference and adjusted for multiple testing. Asterisks denote *FDR* < 10%. **g**, Median number of DEGs and 90% confidence interval (CI) between samples with unmutated and mutated IGHV status for each treatment at 10% *FDR* split into genes with lower or higher expression in U-CLL compared to M-CLL. **h**, Median number of DEGs and 90% CI between samples with and without trisomy 12 for each treatment at 10% *FDR* split into genes with lower or higher expression in trisomy 12 compared to non-trisomy 12, and whether genes are encoded on chromosome 12.

### Pathway dependency of known CLL drivers

We aimed to understand how signalling activities contribute to signatures associated with major CLL subtypes that affect clinical outcome. The proliferative drive of CLL (CLL-PD), linked to prognosis and increased mTOR-MYC-OXPHOS activity, was significantly reduced upon AKT, BET and mTOR inhibition, demonstrating a direct dependency of pro-proliferative capacities of CLL cells on these pathways^12^ (Fig. 3f). The IGHV status is the strongest modulator of the transcriptional variation in CLL and a key determinant of outcome (Fig. 2c, Suppl. Fig. 2f-j). Some well-known markers of IGHV status, i. e. *ZAP70* and *CD38*, were stable after drug treatment, while other signature genes, i. e. *CCL3* and *CD72*, were dependent on BCR activity, suggesting that active signalling maintains part of the IGHV signature (Suppl. Fig. 7a,b). To test the impact of the drugs on the overall signature, we quantified the differentially expressed genes between patients with unmutated and mutated IGHV with each treatment and found that the signature size was moderately reduced by PI3K, mTOR and BTK inhibition (Fig. 3g). This partial reversal was driven by downregulation of IGHV signature genes in unmutated patients which enriched in the cholesterol and glucose metabolism, allograft rejection, adipogenesis, mTORC1 signalling and IFN signalling gene sets (Suppl. Fig. 7c,d). When the BTK and IRAK4 inhibitors were combined, we observed the most pronounced collapse of the IGHV signature, with an enrichment of genes linked to ribosomes and translation, in line with a more pronounced effect on these downstream consequences when the BCR and TLR were simultaneously perturbed^32^ (Fig. 3g, Suppl. Fig. 7c,e). To understand to what extent BCR and TLR signalling cooperate in CLL, we assessed how the sum of the individual BTK and IRAK4 inhibitor effects - with the expectation that there is no interaction - compared to the observed effect from the BTK/IRAK4 inhibitor combination. We found that genes affected by BTK, IRAK4 or BTK/IRAK4 inhibition showed stronger effects than what would be expected from the sum of the individual effects (Suppl. Fig. 7f). Cooperation of TLR and BCR signalling has been described in other B-NHL, where it is linked to activating gene mutations targeting the pathway^33,34^. Our finding suggests a cooperation between TLR and BCR signalling in CLL even in the absence of such a driver mutation.

Trisomy 12 occurs in 15%–20% of CLL patients and is considered an early event^26^. It is associated with disease progression and a higher incidence of aggressive transformation, but its molecular function is not well understood^35,36^. We repeated the same analysis with the trisomy 12 signature. Trisomy 12 signature genes showed partial dependencies on BTK, PI3K and mTOR (Fig. 3h, Suppl. Fig. 7g-i). Differential expression of signature genes encoded on chromosome 12 was largely stable and not affected by disruption of signalling activities, in line with a gene dosage-driven “passenger” signature with no driver role (Fig. 3h). In contrast, we found more genes to discriminate trisomy 12 after MEK, BTK/IRAK4 and MDM2 inhibitor treatment. These findings suggest a modulation of these pathways by trisomy 12 (Fig. 3h, Suppl. Fig. 7j).

### Disease drivers influence pathway connectivity in CLL

Epistasis describes a phenomenon of genetic interactions in which the phenotype of a mutation is modified by the presence or absence of mutations in one or more genes in a non-additive manner^37^. In this study, we adopted this concept and systematically tested for drug-gene interactions in linear models and identified modifications in the transcriptional phenotype of a chemically perturbed gene target in the presence of unmutated IGHV and the common disease drivers trisomy 12, *BRAF*, deletion 17p, deletion 15q and *TP53* (Fig. 4a, Suppl. Fig. 8a). We then grouped the observed phenotypes of the epistatic interactions based on the direction of which perturbations mask, modify, or otherwise influence the expression of genes. We identified genes upregulated by a given aberration which were downregulated in the presence of both perturbation and driver to levels below baseline and vice-versa (inversion; n = 247). Suppression of the driver phenotype in the presence of a perturbation was observed for 112 genes. We also observed synergy (n = 94), where drug effects are amplified and only discernible in the presence of both aberration and drug treatment. Suppression of drug effects in the presence of an aberration was found for 60 genes (Fig. 4b-c, Suppl. Fig. 8b-d). We used these categories to understand the functional implications of disease drivers based on known drug targets or mechanisms. A clear example for loss of function (LOF) in CLL was observed with *TP53* mutations or deletion 17p, which showed impaired transcriptional activation of p53 targets, e. g. *MDM2*, *CDKN1A*, in response to MDM2 inhibitor treatment (≙ suppression of drug effect) (Fig. 4d-f, Suppl. Data Fig. 8d). Consistent with the activation of the MAPK pathway, *BRAF* mutations showed most interactions with MEK inhibition (synergy and driver suppression type (Fig. 4d, Suppl. Fig. 8d)). We also identified so far unknown interactions linking mTOR inhibition to deletion 15q (Fig. 4d, Suppl. Fig. 8d). For unmutated IGHV, we observed interactions with mTOR predominantly of the driver and drug suppression type (Fig. 4d, Suppl. Fig. 8d). We also found important therapeutic targets in lymphoma, e. g. *CD19* and *MS4A1*/CD20, in the synergy group. These findings suggest that expression of these targets are dependent on a certain pathway in molecular subgroups which could serve as a basis for designing new treatment strategies (Fig. 4g,h).

**Fig. 4:**
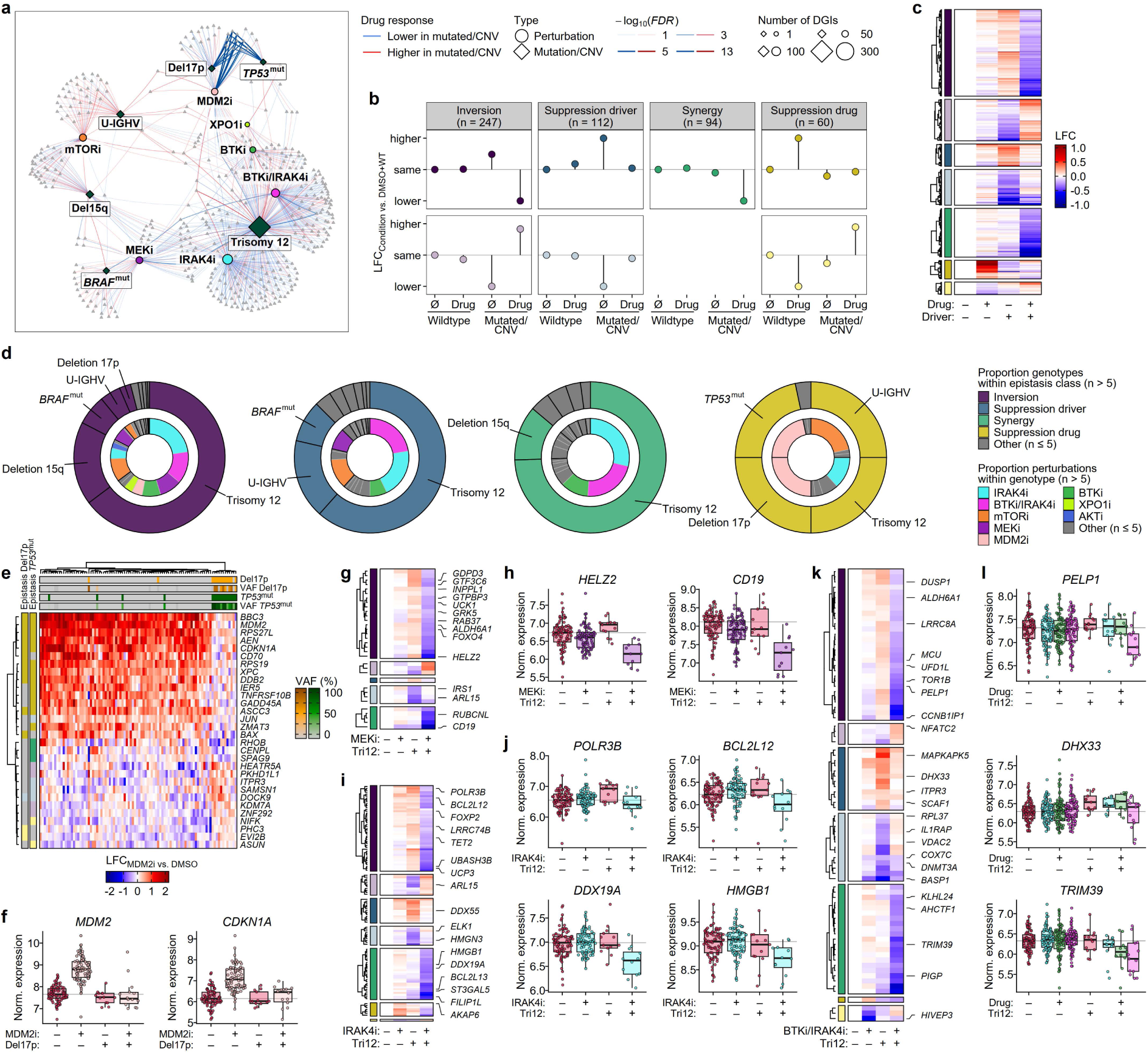
Interaction of perturbation effects and disease drivers. **a**, Pathway connectivity network of significant drug-genotype interactions (DGIs) shown as a bipartite graph. Genes, drugs and disease aberration are connected if a differential transcriptional drug response was observed in the presence of the genetic driver (*FDR* < 10%). Drug-genotype pairs with at least 10 DGIs are displayed. **b**, Schematic classification of epistasis categories based on median values from all genes in the respective groups. **c**, Heatmap of relative gene expression of DGIs in the presence of either drug, genetic driver or both compared to WT/DMSO samples by epistasis class. **d**, Proportion of aberration contributing to each epistasis class (outer circle) and proportion of perturbations contributing to each aberration within each epistasis class (inner circle) with more than 5 DGIs. **e**, Heatmap visualising the effect sizes of genes with significant interaction between MDM2 inhibition and del17p or *TP53*^mut^ (*FDR* < 10%). **f**, Effect of MDM2 inhibition on normalised expression of *MDM2* and *CDKN1A* by deletion 17p status. **g**, Heatmap visualising the effect sizes of genes with significant interaction between MEK inhibition and trisomy 12 (*FDR* < 10%). **h**, Normalised expression of MEK inhibition on *HELZ2* and *CD19* by trisomy 12 status. **i**, Heatmap visualising the effect sizes of genes with significant interaction between IRAK4 inhibition and trisomy 12 (*FDR* < 10%). **j**, Effect of IRAK4 inhibition on normalised expression of *POLR3B*, *BCL2L12*, *DDX19A* and *HMGB1* by trisomy 12 status. **k**, Heatmap visualising the effect sizes of genes with significant interaction between combined BTK/IRAK4 inhibition and trisomy 12 (*FDR* < 10%). **l**, Effect of DMSO (red), IRAK4 (cyan), BTK (green) and BTK/IRAK4 inhibition (pink) on normalised expression of *PELP1*, *DHX33* and *TRIM39* by trisomy 12 status.

Most drug-gene interactions were observed for trisomy 12 (n = 314) characterised by inversion and synergy with BTK, MEK, IRAK4 and combined BTK/IRAK4 inhibition (Fig. 4d,i-l, Suppl. Fig. 8d). The effect of IRAK4 inhibition in the presence of trisomy 12 is remarkable as the inhibitor’s effect was undetectable when the complete cohort was analysed (Fig. 2g). We identified most drug-gene interactions were modulated by trisomy 12 with IRAK4 (n = 114), combined BTK/IRAK4 (n = 83) and BTK inhibition (n = 39) providing further evidence for crosstalk and cooperative effects between BCR and TLR signalling in trisomy 12 (Fig. 4d,i-l). To demonstrate the role of IRAK4 as a mediator of TLR signalling in CLL, we treated patient samples with either IRAK4 inhibitor alone or in combination with the TLR9 agonist CpG. TLR9 stimulation led to increased viability in line with a pro-tumourigenic role of TLR signalling, which was abrogated by IRAK4 inhibition (Suppl. Fig. 9a). We searched for focal gains in the CNV profiles of our CLL cohort (n = 366) which may allow restricting the number of candidate genes targeted by trisomy 12, and identified a CLL case with a focal gain between chr12:42,538,673 and chr12: 44,229,999. This gain included nine protein-coding genes including *IRAK4* (Fig. 5a-c, Suppl. Fig. 9b). We clustered gene expression of CLL samples (n = 210) based on the trisomy 12 baseline signature and restricted the analysis to genes not encoded on chromosome 12 (642 out of 917 total trisomy 12 signature genes) to prevent clustering due to gene dosage. The sample with the focal gain of *IRAK4* clustered with the trisomy 12 samples demonstrating the phenotype resemblance (Fig. 5c).

**Fig. 5:**
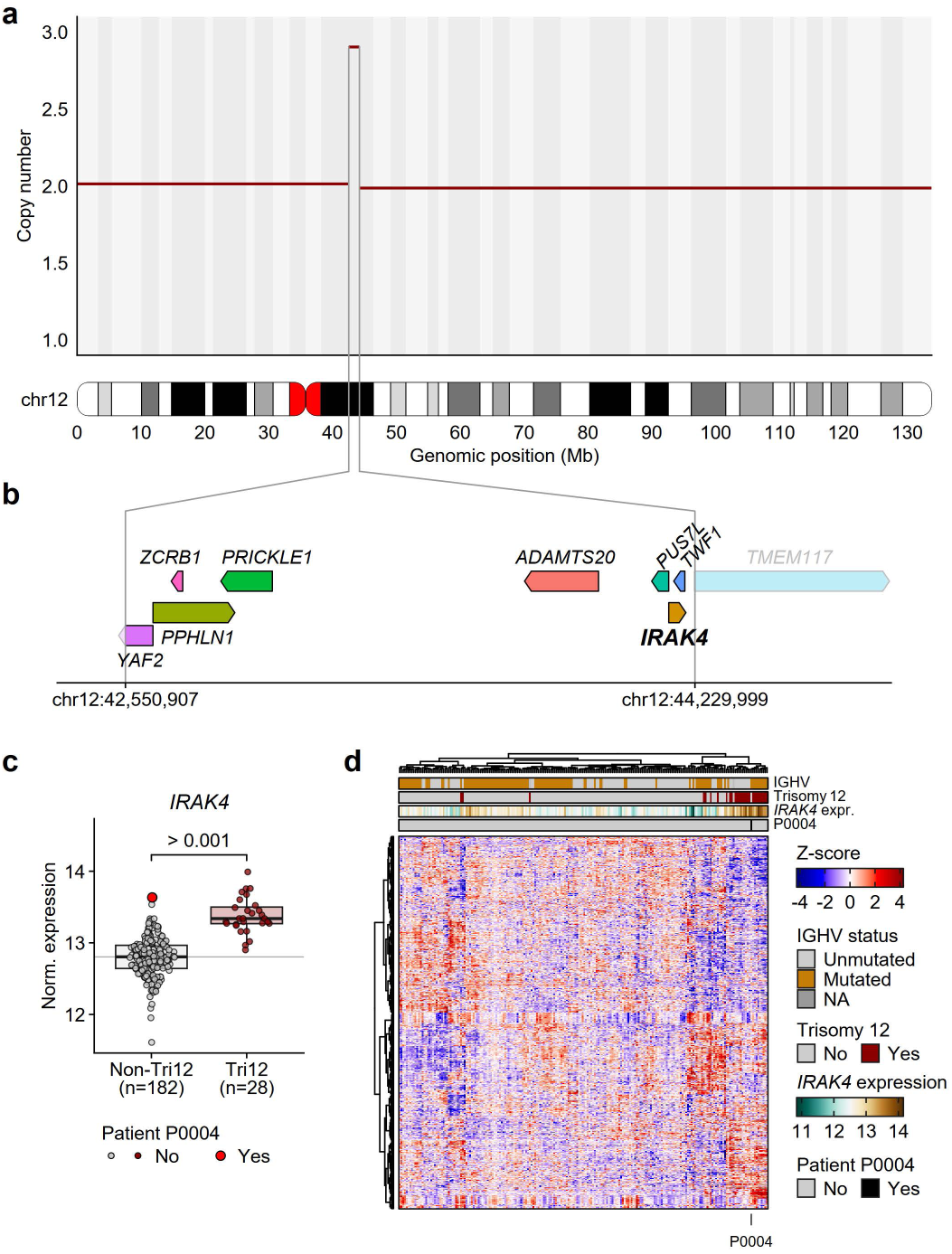
IRAK4 is a disease driver targeted by trisomy 12. **a**, CNV plot with ideogram of chromosome 12 for patient P0004. The red line shows the copy numbers across the chromosome. **b**, Genomic track displaying the protein-coding genes on the amplified locus chr12:42,550,907-chr12: 44,229,999 (hg19). **c**, Gene expression of IRAK4 by trisomy 12 status for the CLL cohort from Lütge et al.^13^. The significance level was calculated using two-tailed Student’s *t*-test and shown as nominal *P*-value. **d**, Heatmap visualising the scaled gene expression of trisomy 12 signature genes excluding the genes on chromosome 12 (n = 642) for the CLL cohort (n = 210) from Lütge et al.^13^

Our findings suggest that trisomy 12 confers increased TLR signalling capacity driven by the increased *IRAK4* expression from gene dosage and cooperates with BCR signalling.

## Discussion

We performed a functional perturbation screening directly in cancer cells at scale to capture drug-induced global gene expression changes with low-depth RNA sequencing. Prior studies investigating perturbation-induced gene expression changes have exploited single-cell technology (few patients, many cells e. g. MIX-seq^38^, RASL-seq^39^ or DRUG-seq^40^) and focused on establishment of technical frameworks, but less on exploiting the signatures to dissect disease biology and cellular processes. The Connectivity Map focused on the differences and signatures induced by a large number of drugs in fewer cell lines^18,19^. By design, it therefore has unique resolution on the drug effects and contributes to the understanding how drugs work. Our approach complements the Connectivity Map by leveraging many primary tumours from an individual disease, and resolving the effects of a limited number of clinically used drugs on a broad and representative sampling of molecularly diverse tumours without immortalisation artefacts. We exploit the drug effects on transcriptomes to resolve disease mechanisms as well as inter-patient heterogeneity. This design allowed us to measure downstream effects of signalling events even if effects are subtle. Our findings suggest that insights and hypotheses from gene expression studies on primary tumours may be vastly increased by the analysis of such perturbed samples as the ability to specifically target molecules and pathways grows. Multiple clinical inhibitors targeting PI3K, BTK, mTOR and MEK, were used in this study as optimal combination and clinical assignment to patients is a challenge, which cannot be easily addressed in trials^41^.

We used interaction testing to study the effect of gene mutations on the perturbed gene expression, e.g. by LOF mutations (*TP53*) or GOF (trisomy 12, *BRAF*). A particularly notable example is trisomy 12, present in approximately 20% of CLL patients. We identified IRAK4 as a driver in CLL with trisomy 12, highlighting how perturbation transcriptomics can uncover actionable targets within genetically defined subgroups. In line with our proposed mechanism where IRAK4 is a key target, trisomy 12 has been found to be strongly associated with response to TLR stimuli such as resiquimod (TLR7/8) and CpG-ODN (TLR9)^42^. Reid et al. studied genes targeted by trisomy 12 in a human pluripotent cell line model and identified IRAK4 inhibition to be potent against human trisomy 12-harboring CLL cells in patient-derived xenografts^43^.

Our experiments enable the deconvolution of drug effects by factor analysis, and facilitate the transition to patient-specific pathway assembly read-outs by drug perturbation and global expression analysis (Fig. 6). The identification of compensatory pathways (factor 14, 20) may provide a basis for optimised combination treatment. While prior drug response perturbation studies have hinted at some of these findings and structure^11^, the use of a large set of tumours and high-dimensional readout of gene expression gives a more comprehensive and systematic picture of the connectivity of signalling pathways in CLL, where effects of inhibitors on individual pathways can be deconvoluted and linked to other drugs. On the route to improved precision medicine for individual cancer patients, the workflows established in our study provide a framework to measure pathway connectivity of primary cancer cells using clinically relevant drugs and adapting the treatment along the lines of measured pathway activity which can be generated by the projection of the individual patients response compared to a larger reference set of measurements such as the one presented here. While the precise strategy for optimal drug combinations will need to be demonstrated, our data provides a roadmap for such experimentation.

**Fig. 6:**
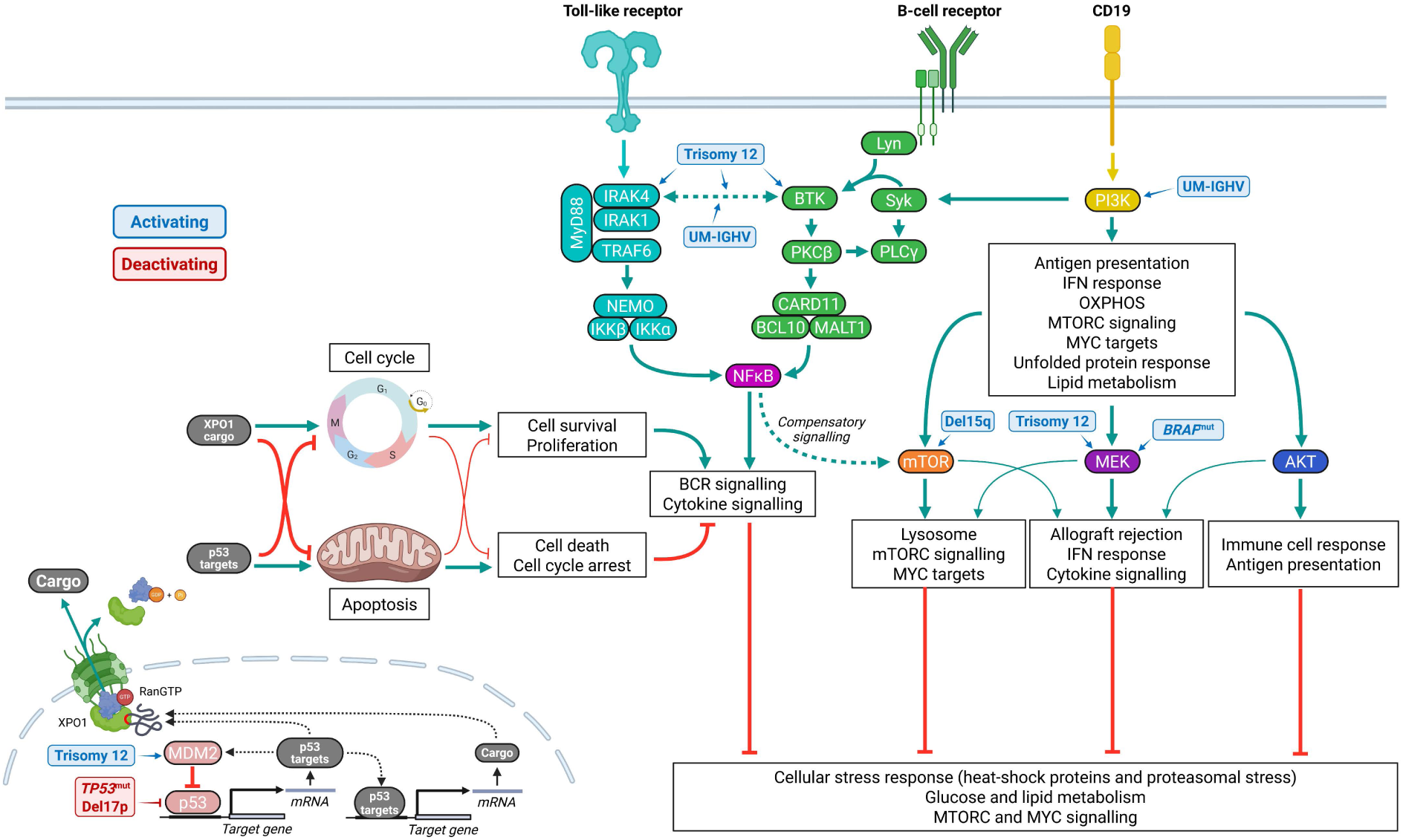
Pathway diagram and dependency map in CLL. Pathway dependencies modulated by gene mutations/CNVs or IGHV status were identified by drug-genotype interaction analysis and depicted as activating (blue) or inhibitory (red) (https://BioRender.com/svmtjkd).

Drug-perturbed transcriptome profiling measures connectivity of cancer pathway dependencies and functionally annotates mutations. As more specific drugs become available, tumour pathway wiring could be fully resolved and improve precision oncology, moving closer to treatments guided by measured pathway activity rather than inferred potential.

## Material and Methods

### Patient samples and study approval

Patient material was acquired in accordance with the Declaration of Helsinki and approval from the local ethics committees (S-206/2011; S-356/2013, 2009-0062; 2019/01744), after written informed consent. Detailed patient information is listed in Suppl. Table 1. DNA mutations, CNVs, DNA methylation and gene expression were performed as described previously^11^.

### Isolation and cryoconservation of MNCs from whole blood

Mononuclear cells from peripheral blood were isolated using Ficoll-Paque (Cytiva) density gradient centrifugation, cryopreserved in freezing medium (72.5% RPMI-1640 (Gibco), 20% fetal bovine serum (FBS, Gibco), 7.5% DMSO (Sigma-Adrich)) and stored at −180°C in liquid nitrogen.

### Compounds

Small-molecules (except compound 26 (Calbiochem)) were purchased from Selleckchem and reconstituted in DMSO at 10 mM.

In a pilot, we treated four CLL samples with concentrations based on results from multiple prior drug screens^11,12^ with average toxicity below 20% (Suppl. Methods Fig. 1a,b) and confirmed drug-specific gene expression changes in line with the drugs’ mechanisms of action (Suppl. Methods Fig. 1c-h). For the screen, stocks were diluted to 200-fold of the final assay concentrations with DMSO as “ready-to-use” aliquots and stored at −20°C.

### *Ex-vivo* drug treatment and cell viability assessment

Primary samples were thawed and incubated in RPMI-1640 supplemented with 10% heat-inactivated human serum from male AB clotted whole blood (Sigma-Aldrich), 1% 10,000 U/mL penicillin/10,000 µg/mL streptomycin (P/S, Gibco) and 2% 200 mM L-glutamine (Gln, Gibco) (RPMI-HS) on a roller mixer for 3h at room temperature to allow recovery from the thawing process and support the removal of DMSO from the freezing medium. The medium was replaced and 5 x 10^6^ cells were treated with each drug (Suppl. Methods Fig. 1c) and cultured at 37°C and 5% CO_2_ for 48h. Cells were harvested, washed twice with cold phosphate-buffered saline (Gibco), lysed in 600 µL TRI Reagent (Zymo Research) and stored at −80°C until RNA extraction.

Sample viabilities of each sample (before and after perturbation) were assessed on the flow cytometer LSR II Fortessa with FACSDiva v8.0.1 software (BD Biosciences) and analysed with FlowJo v10.10.0 (FlowJo LLC) (Suppl. Method Fig. 2).

### TLR9 stimulation and ATP-based viability assessment

For the 5x concentrated working dilution, a 5-point 5-fold serial dilution of IRAK4i was prepared with a starting concentration of 50 µM (≙ 10 µM–0.016 nM final assay concentration). For the negative control, 5 µL of 0.5% DMSO working dilution was pre-plated (≙ 0.1% final concentration).

MNCs from 20 CLL patients were thawed and incubated for 3h on the roller mixer prior to seeding as described above and reconstituted to 1 x 10^6^/mL (1.25X) cell concentration. In unstimulated conditions, 20 µL (≙ 20,000 cells) were added to each well containing the pre-plated DMSO or IRAK4i. For the CpG stimulation, cells were supplemented with 1.25 µg/mL of the TLR9 agonist ODN2006 (InvivoGen) before seeding to reach a final assay concentration of 1 µg/mL. Cells were incubated at 37°C and 5% CO_2_ for 48h. For the ATP-based viability assessment, plates were spun down briefly before adding 7 µL CellTiter-Glo (Promega) to each well. The reaction was incubated for 20 min at RT and luminescence measured on a BioTek Synergy LX multimode microplate reader (Agilent). To obtain normalised viabilities, intensity values were divided by the median of all DMSO values for each condition and concentration for each patient.

### RNA extraction

Frozen lysates were thawed at 37°C for 3 min. RNA was precipitated by thoroughly mixing 100 µL of 100% ethanol (Sigma-Aldrich) with the lysate. RNA was extracted with the Direct-zol RNA MiniPrep kit (Zymo Research) including on-column DNase treatment (30U per sample) for 15 min at room temperature according to the manufacturer’s protocol and eluted in 25 µL nuclease-free water (QIAGEN). The concentration was determined with Quant-it RiboGreen RNA assay (ThermoFisher Scientific) on a BioTek Synergy LX multimode microplate reader. RNA was stored at −80°C in 5 µL nuclease-free water until use for library preparation and RNA sequencing.

### Library preparation and 3’ RNA sequencing

For the pilot screen (4 patients) 200 ng of total RNA were used for cDNA library generation with the QuantSeq 3′ mRNA-Seq Library Prep Kit FWD for Illumina (Lexogen). Libraries from 42 samples were pooled and sequenced on the NextSeq500 (Illumina) using 75 bp and single-end reads yielding 3.9 to 13.5 million reads. For the comparison between the library preparation methods, baseline profiles of 5 patients generated by whole-transcript deep sequencing (TruSeq, Illumina)^13^, the same RNA material was sequenced in the pilot screen.

For the remaining 112 patients, cDNA libraries were prepared from 100 ng total RNA with the QuantSeq 3′ mRNA-Seq Library Prep Kit FWD for Illumina (Lexogen). Up to 48 libraries (22-48) were pooled and sequenced on the HiSeq 4000 (Illumina) using 50 bp and single-end reads yielding 2.3 to 21.7 million reads.

### Processing of the 3’-RNA sequencing data

The same libraries from the first sequencing run were sequenced twice and the resulting reads were pooled. The demultiplexed RNA sequencing FASTQ files were mapped to the ENSEMBL human reference genome release 104 (Homo sapiens GRCh38.104) using STAR (version 2.7.9a) with default parameters. The aligned reads were summarised into per-gene counts using htseq-count (version 0.9.1) with default parameters in union mode, counting only those reads unambiguously mapped to a single gene. The resulting gene count matrix was imported into R (version 4.5.1) for further processing. Given the shallow sequencing depth, the gene count matrix exhibited higher sparsity compared to traditional bulk RNA-seq data. This sparsity posed challenges for accurately estimating between-sample normalisation factors using methods designed for deep RNA sequencing, such as DESeq2^44^. To address this, scTransform^45^, a method that applies regularised negative binomial regression to sparse gene count matrices from single-cell sequencing data, was used to estimate the normalisation factors. These factors were then supplied to DESeq2 (version 1.44) for variance-stabilising transformation and differential expression analysis. For all the downstream analyses, only protein-coding genes with minimum 10 counts in at least one sample were retained.

The baseline profiles were combined with profiles generated by 3’-end shallow depth sequencing (QuantSeq, Lexogen) and normalised for library size before comparative analysis.

### Data analysis

#### Statistics

Correlation trends were indicated by Pearson correlation (*R*) and linear regression lines with 95% confidence intervals. Density distributions of single values were visualised with sina plots and summarised with Tukey-style box and whiskers plots. Statistical differences between groups were computed with unpaired two-tailed Student’s *t*-test when groups contained samples from different patients, e. g. mutant vs. wild-type. Paired two-tailed Student’s *t*-tests were used when conditions of the same patients were compared. Nominal *P*-values were corrected for multiple testing using the Benjamini-Hochberg procedure and indicated as false-discovery rate (*FDR*).

#### Dimensionality reduction and clustering analysis

Dimensionality reduction including principal component analysis (PCA), *t*-distributed Stochastic Neighbor Embedding (*t*-SNE) and hierarchical clustering were conducted on gene counts after variance-stabilising transformation (*vst* function from DESeq2 package). PCA (*prcomp* function from base R) and *t*-SNE (*tsne* function from M3C package^46^, version 1.26) were performed on the 1,000 most variable genes, unless indicated. Hierarchical clustering was carried out using the Ward.D2 method based on Euclidean distances computed from the centered and scaled expression values of genes selected from differential expression analyses. The selection criteria are detailed in the figure legends. The ComplexHeatmap package^47^ (version 2.20) was used for visualisation. To analyse expression patterns after drug treatment, patient-specific variations were adjusted in the gene expression matrix before dimension reduction (*removeBatchEffect* function from limma package^48^, version 3.60.6). The application of this adjustment is noted in the respective figure legends.

#### Differential expression analysis

All differential expression analyses were conducted on the scTransform-adjusted gene count matrix using the R package DESeq2^44^ (versions 1.44.0 and 1.48.2).

To compute genes differentially expressed between drug- and DMSO-treated samples, patient IDs were included as a covariate in the design formula to account for patient-specific variation. Log2-fold changes were shrunken with the *lfcShrink* function using the apeglm^49^ method. Genes were filtered by significance (*FDR* < 0.5%) and effect size (|fold change| > 1.05). p53 expression scores were downloaded from the Target Gene Regulation Database 2.0^50^. The CLL-PD score was computed from the Z-score gene count matrix using *CLLPDestimate* from the mofaCLL package^12^ (version 0.0.9).

The transcriptional signature of a disease dimension was defined as the set of differentially expressed genes between molecular subgroups individually for each perturbation. To account for the confounding effects of IGHV and trisomy 12 status, these variables were included in the design formula. To compare the number of signature genes between perturbations, the 90% confidence interval was estimated using bootstrapping (1,000 times, sampling 90% of patients while maintaining the ratio between wild-type and mutated/CNV patients).

#### Drug-genotype interaction testing and epistasis clusters

The interactions between genetic aberrations and drug effects were modeled by linear interaction testing with DESeq2 using a design formula that accounts for aberration, patient ID and treatment condition. Drug-gene interaction genes were filtered by *FDR* (< 10%). Median and scaled expression was calculated for each gene and group (DMSO/wild-type, drug/wild-type, DMSO/mut, drug/mut) and normalised against DMSO/wild-type as the reference. Interaction types were grouped using soft Fuzzy C-Means Clustering (FCM) with the *cmeans* function (centers = 20, iter.max = 1000, dist = “euclidian”, method = “cmeans”, m = 1.4) from the e1071 package^51^ (version 1.7.16). FCM was repeated three times for genes with probability scores below 0.85 until every gene was assigned to a cluster with a probability score of at least 0.85, resulting in 60 pre-clusters. Pre-clusters with similar patterns were summarised to 20 final clusters using harder FCM (m = 1.1) and manual curation.

#### Pathway connectivity networks

Pathway connectivity networks of significant DEGs are shown as bipartite graphs in which perturbation and DEGs denote the disjoint and independent sets of nodes and are connected by the edges. The edges are weighted by the significance level of the drug effects. For the connectivity network of interactions, nodes were filtered for aberrations with more than 5 mutated/CNV cases. The edges are weighted by the significance level of the interaction. Node coordinates were computed using the network package^52^ (version 1.19.0).

#### Guided sparse factor analysis

Guided Sparse Factor Analysis (GSFA) was conducted using the GSFA package (version 0.2.8)^31^. The gene count matrix was pre-processed by accounting for the patient-specific variation and selecting the 5,000 genes with the largest deviance after deviance residual transformation^53^. DMSO-treated samples were indicated as negative controls, and a “mixture_normal” distribution was used as the sparse prior. Singular value decomposition was employed to initialise the factor values, followed by 3,000 iterations of Gibbs sampling to train the model. Posterior means were calculated using the last 1,000 iterations. All other parameters were used with default settings.

The model with 20 factors was chosen for further analysis (Suppl. Method Fig. 3). Perturbations and genes with association effect sizes above 0.25 and posterior inclusion probabilities (*PIP*) above 0.95 were considered to have a significant effect on a particular factor. Multiple linear regression was conducted with Lasso penalisation, leave-one-out cross-validation, mean squared error loss, 200 iterations, mutations/CNVs as predictors and centered and scaled individual drug effects as response vectors using the glmnet package^54^ (version 4.1-8).

#### Pathway enrichment analysis

Parametric analysis of gene set enrichment^55^ was performed on the DEGs (ranked by *t*-statistics) or genes associated with GSFA factors (ranked by factor loading). Human gene set collections (version 2024.1) were downloaded from MSigDB (https://www.gsea-msigdb.org/gsea/msigdb/human/collections.jsp). The decoupleR package (version 2.14.0) was used to infer the impact on transcription factor activity by each drug treatment^28^. The filter criteria for model input and results are indicated in the respective figure legends.

## Data availability

All RNA sequencing data generated in this study will be available at European Genome-Phenome Archive (EGA) upon publication. The sequencing data generated in our previous studies and used in the current study, including whole-exome/whole-genome sequencing, targeted-sequencing, DNA methylation profiling and RNA sequencing of baseline patient samples, are available at EGA under the accession code EGAS00001001746. Processed data, including RNA sequencing data, drug sensitivity profiling and sample metadata are available in our project repository at GitHub (https://github.com/THLDo/CLL_Connectivity). The RNA drug perturbation dataset can also be explored interactively via the Shiny app available upon publication.

## Code availability

The computational codes for reproducing all figures and results reported in this article will be provided in our project repository at GitHub (https://github.com/THLDo/CLL_Connectivity) under the GNU General Public License v3.0 upon publication.

## Acknowledgements

T. Zenz was supported by the CRPP ‘Next Generation Drug Response Profiling for Personalized Cancer Care’, the Swiss Cancer Research foundation (KFS-4439-02-2018), the Monique-Dornonville-de-la-Cour Stiftung and the Helmut Horten Stiftung. M. F. Pohly, W. Huber were supported by the INTeRCePT, and T. Zenz by the INTeRCePT and POLAR (Uniscientia Stiftung) projects funded by The LOOP Zurich. J. Lu, C. Lohoff and W. Huber were supported by the SMART-CARE project funded by the Federal Ministry of Research, Technology and Space of Germany (161L0213 and 16LW0237). For access to infrastructure, we thank M. Bawohl, M. Rechsteiner and A. Weber at the Department of Pathology and Molecular Pathology, and the Functional Genomics Center Zurich. For access to the EMBL Heidelberg HPC Cluster, we gratefully thank J. Pečar and EMBL IT Services.

## Contributions

THLD, JL, WH, TZ conceptualised the study and designed the experiments. THLD, SK, MFP and FJ conducted experiments. SS, JH, WH, JL and TZ were responsible for data curation. CL, FJ and VB performed RNA sequencing and data pre-processing. THLD, CL, JL and TZ performed data analysis and data visualisation. THLD, CL, JL, WH and TZ interpreted the data. THLD, CL, JL, WH and TZ wrote the original draft of the manuscript. All authors reviewed and edited the manuscript.

## Supplementary figures

**Suppl. Fig. 1:**
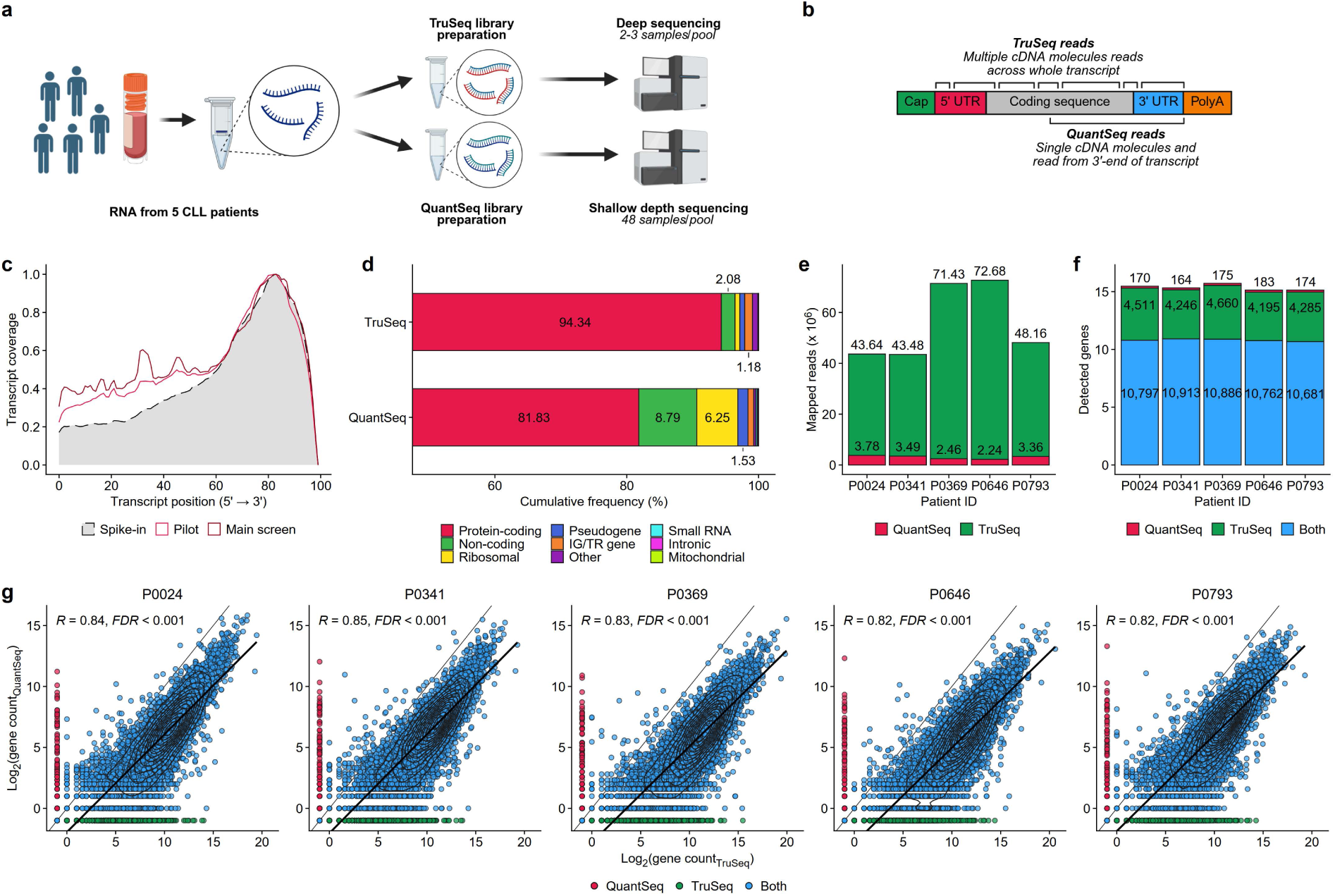
Comparison of shallow 3’-QuantSeq to regular coverage whole-transcript TruSeq sequencing. **a**, Overview of experimental workflow (https://BioRender.com/u4e6koa). **b**, Schematic comparison of read distribution along transcript between 3’-QuantSeq^23^ and whole-transcript TruSeq (https://BioRender.com/sx3vfjw). **c**, Biased read coverage of transcript 3’-end for exemplary samples. **d**, Mapped biotypes, **e**, number of mapped reads, **f**, and number of detected genes by method. Colours depict whether genes were detected by both methods, only by TruSeq or only by QuantSeq. **g**, Correlation of log2 gene expression counts detected by both methods.

**Suppl. Fig. 2:**
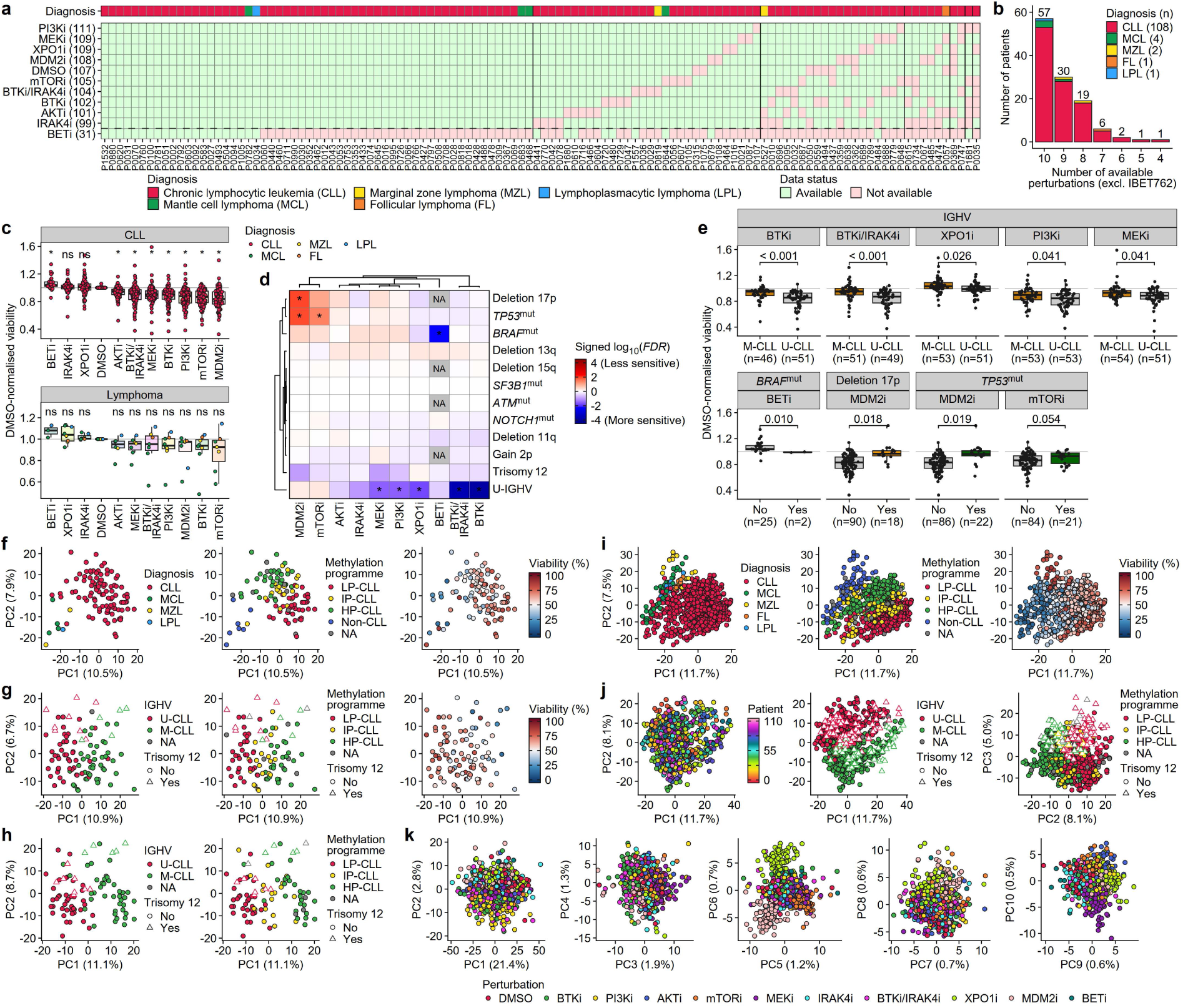
Overview viability effect of drugs and transcriptional landscape. **a**, Overview of completeness and **b**, number of available treatment conditions by patient. **c**, DMSO-normalised sample viability after 48h treatment measured by flow cytometry and separated by diagnosis. Significance levels were calculated using two-tailed paired Student’s *t*-test and adjusted for multiple testing. Asterisk denotes significant differences between drug-and DMSO-treated samples (*FDR* < 10%). **d**, Genomic association with drug response for CLL samples. Asterisk denotes associations with significant differences in drug response between patients with and without respective genetic aberrations with more than 5 cases (*FDR* < 10%). Significance levels were calculated using two-tailed Student’s *t*-test and adjusted for multiple testing. **e**, Boxplots of individual patient data shown for top significant associations (*FDR* < 10%). Significance levels were calculated using two-tailed Student’s *t*-test and adjusted for multiple testing. **f**, PCA based on 1,000 most variant genes from DMSO-treated CLL and lymphoma samples (n = 107), **g**, only CLL samples (n = 100), and **h**, from overlapping CLL patients from Lütge et al.^13^ profiled by deep whole-transcript RNA sequencing (n = 95). **i**, PCA based on 1,000 most variant genes from DMSO- and drug-treated CLL and lymphoma samples (n = 1,086), **j**, only CLL samples (n = 1,010), **k**, and after adjustment for the patient effect (n = 1,010).

**Suppl. Fig. 3:**
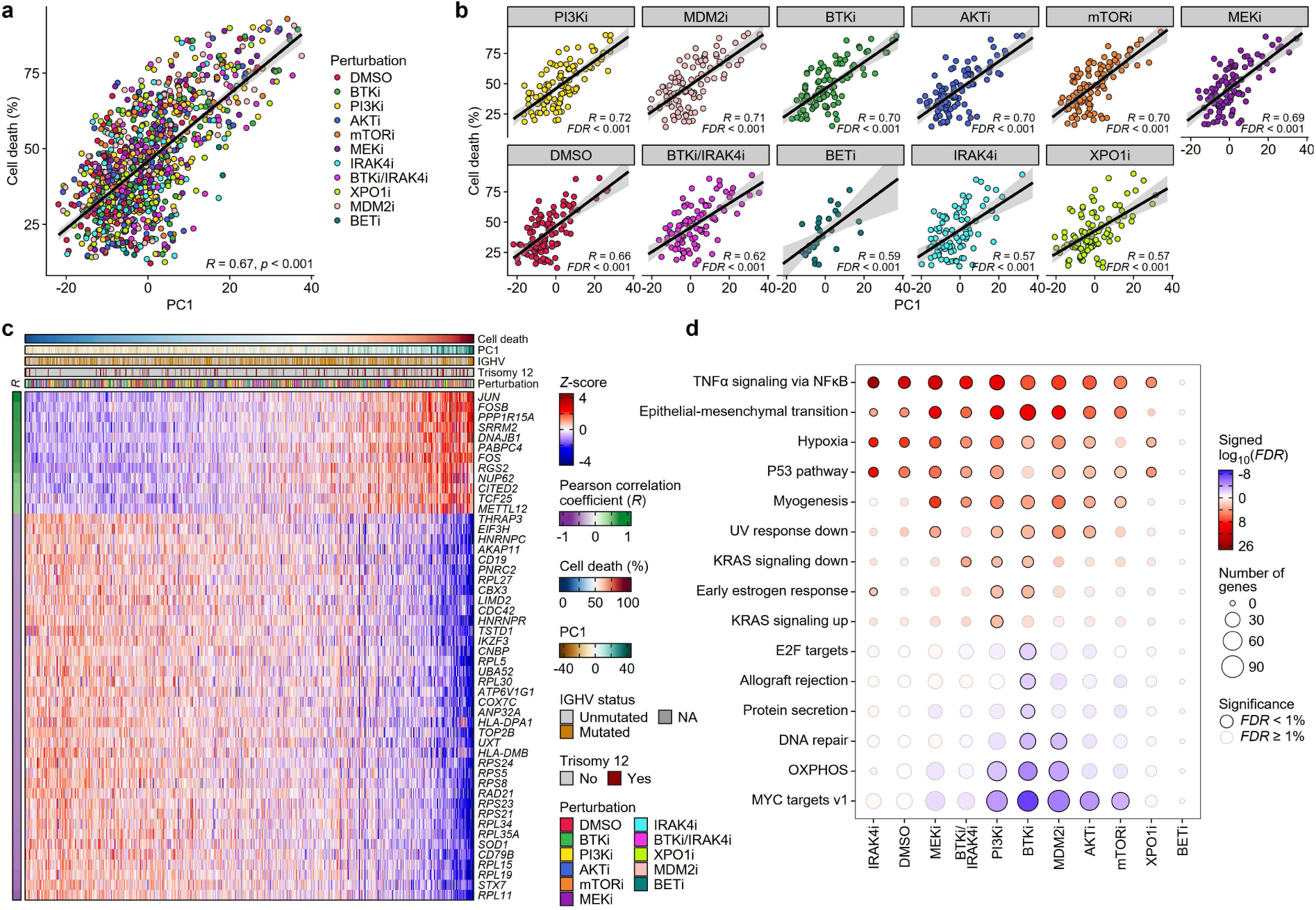
Transcriptional signature associated with viability. **a**, Correlation of cell death (100% - sample viability) with PC1 for all samples and **b**, by treatment. Correlation trends are indicated by Pearson correlation coefficient (*R*) and adjusted for multiple testing. **c**, Heatmap showing normalised expression of top 50 correlated genes with cell death. Correlation trends are indicated by Pearson correlation coefficient (*R*). **d**, GSEA individually for each treatment using genes with significant correlations between expression and cell death (*FDR* < 0.5%) using hallmark gene sets and number of enriched genes (< 10). Gene sets were filtered by *FDR* (< 1%). Enrichments indicate positive (red) or negative (blue) associations with cell death.

**Suppl. Fig. 4:**
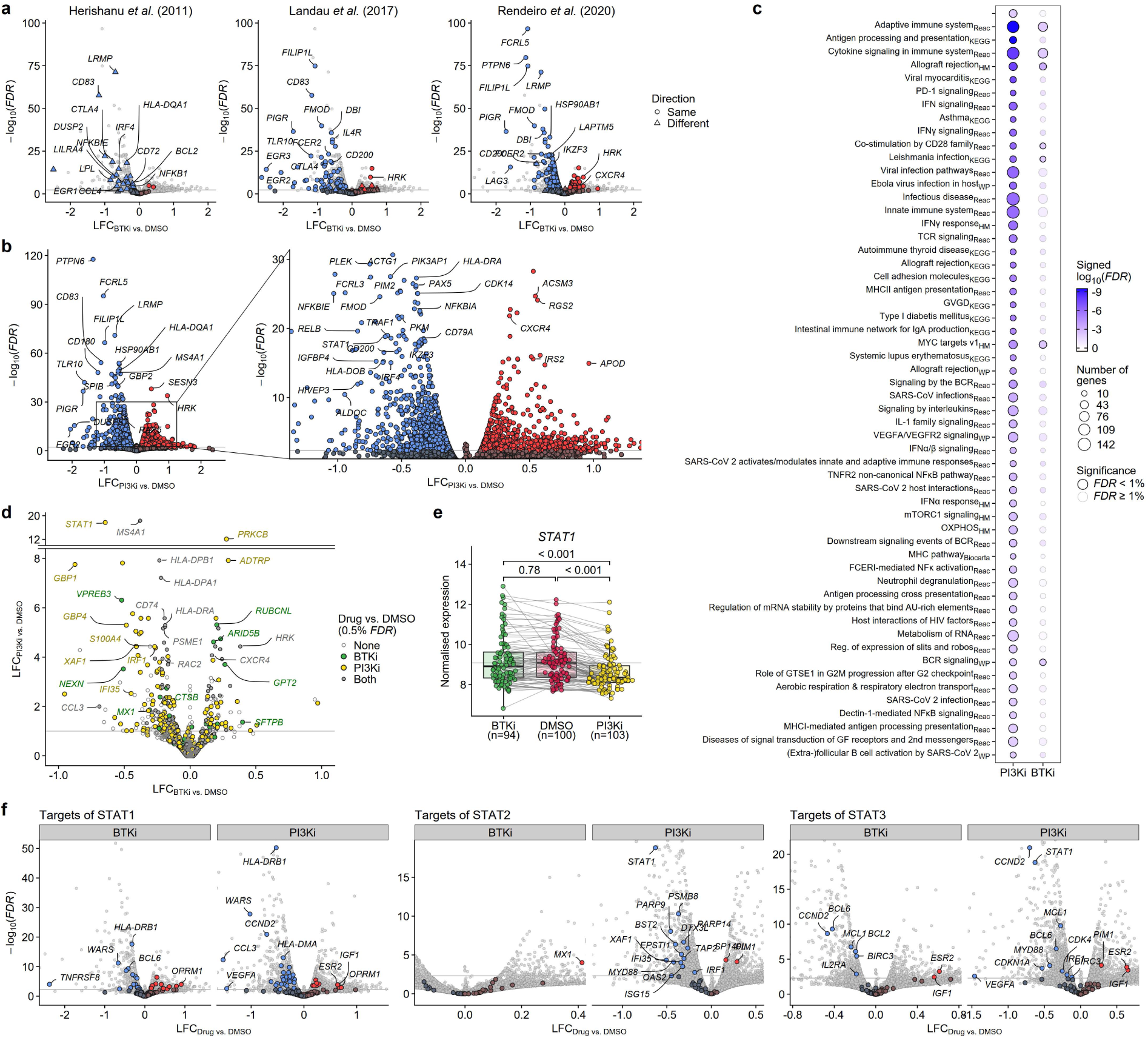
Drug effects of BTK and PI3K inhibitors on BCR inhibition. **a**, Projection of data from Herishanu et al.^25^ (*ex-vivo* BCR activation signature), Landau et al.^56^ and Rendeiro et al.^27^ *(in-vivo* BTK inhibitor signature) into volcano plot of differential gene expression after BTK inhibition. Shapes depict whether the gene is regulated into the same or different direction. **b**, Volcano plot summarising the differential gene expression after PI3K inhibition. **c**, GSEA with DEGs after BTK and PI3K inhibitor treatment (0.5% *FDR*) using hallmark and canonical pathways. Gene sets were then filtered by *FDR* (< 1%) and number of enriched genes (> 10). Depicted are all significant gene sets for PI3K inhibition. **d**, Volcano plot summarising the differential gene expression between PI3K and BTK inhibition. Colours and shapes depict in which comparisons the gene was deregulated (0.5% *FDR*). **e**, Normalised *STAT1* expression in samples treated with DMSO, BTK or PI3K inhibitor. Significance levels were calculated using two-tailed paired Student’s *t*-test and shown as nominal *P*-values. **f**, Projection of STAT1, STAT2 and STAT3 transcription factor target genes into volcano plot of differential expression after BTK and PI3K inhibition.

**Suppl. Fig. 5:**
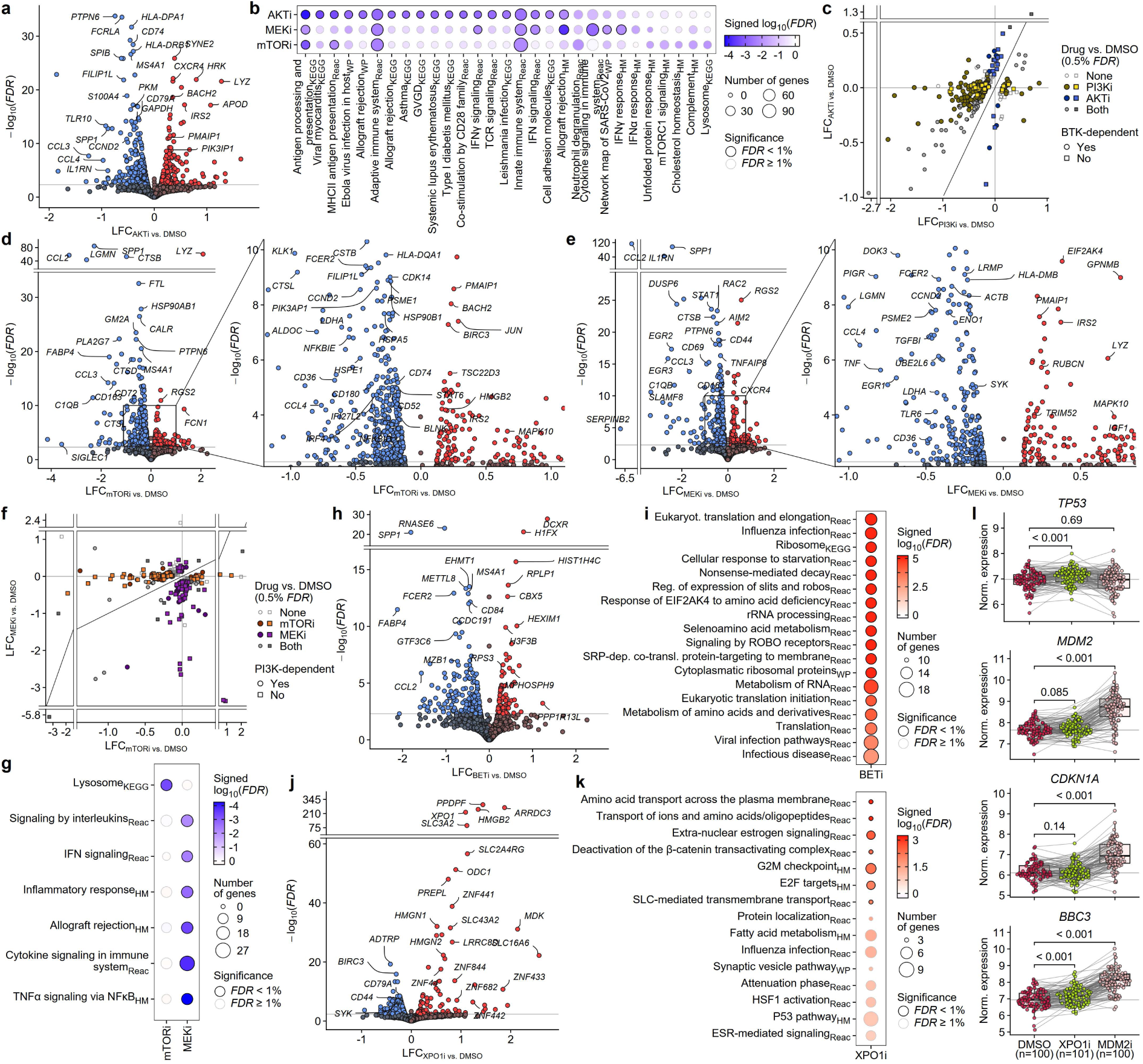
Drug effects of AKT, mTOR, MEK, XPO1 and BET inhibitors. **a**, Volcano plot summarising gene expression after AKT inhibitor treatment. **b**, GSEA with DEGs after AKT, mTOR and MEK inhibitor treatment (0.5% *FDR*) using hallmark and canonical pathways gene sets. Gene sets were filtered by *FDR* (< 1%) and number of enriched genes (> 10). Depicted are all significant gene sets for AKT and the top 10 gene sets for MEK and mTOR inhibition. **c**, Scatter plot of drug effect sizes for DEGs between AKT and PI3K inhibitor treatment (0.5% *FDR*). Colours and shapes depict in which comparisons the gene was deregulated (0.5% *FDR*). **d**, Volcano plot summarising gene expression after mTOR and **e**, MEK inhibitor treatment. **f**, Scatter plot of drug effect sizes for DEGs between MEK and mTOR inhibitor treatment (0.5% *FDR*). Colours and shapes depict in which comparisons the gene was deregulated (0.5% FDR). **g**, GSEA with genes differentially expressed between MEK and mTOR inhibition (0.5% *FDR*) using hallmark and canonical pathways gene sets. Gene sets were filtered by *FDR* (< 1%) and number of enriched genes (> 10). **h**, Volcano plot summarising gene expression after BET inhibitor treatment. **i**, GSEA with DEGs after BET inhibitor treatment (0.5% *FDR*) using hallmark and canonical pathways gene sets. Gene sets were filtered by *FDR* (< 1%) and number of enriched genes (> 10). **j**, Volcano plot summarising gene expression after XPO inhibitor treatment. **k**, GSEA with DEGs after XPO1 inhibitor treatment (0.5% *FDR*) using hallmark and canonical pathways gene sets. Gene sets were filtered by *FDR* (< 1%) and number of enriched genes (≥ 3). **l**, Effect of XPO1 (green) and MDM2 inhibition (rose) on the expression of *TP53* and the p53 target genes *MDM2*, *CDKN1A*, *BBC3*. Significance levels were calculated using two-tailed paired Student’s *t*-test and shown as nominal *P*-values.

**Suppl. Fig. 6:**
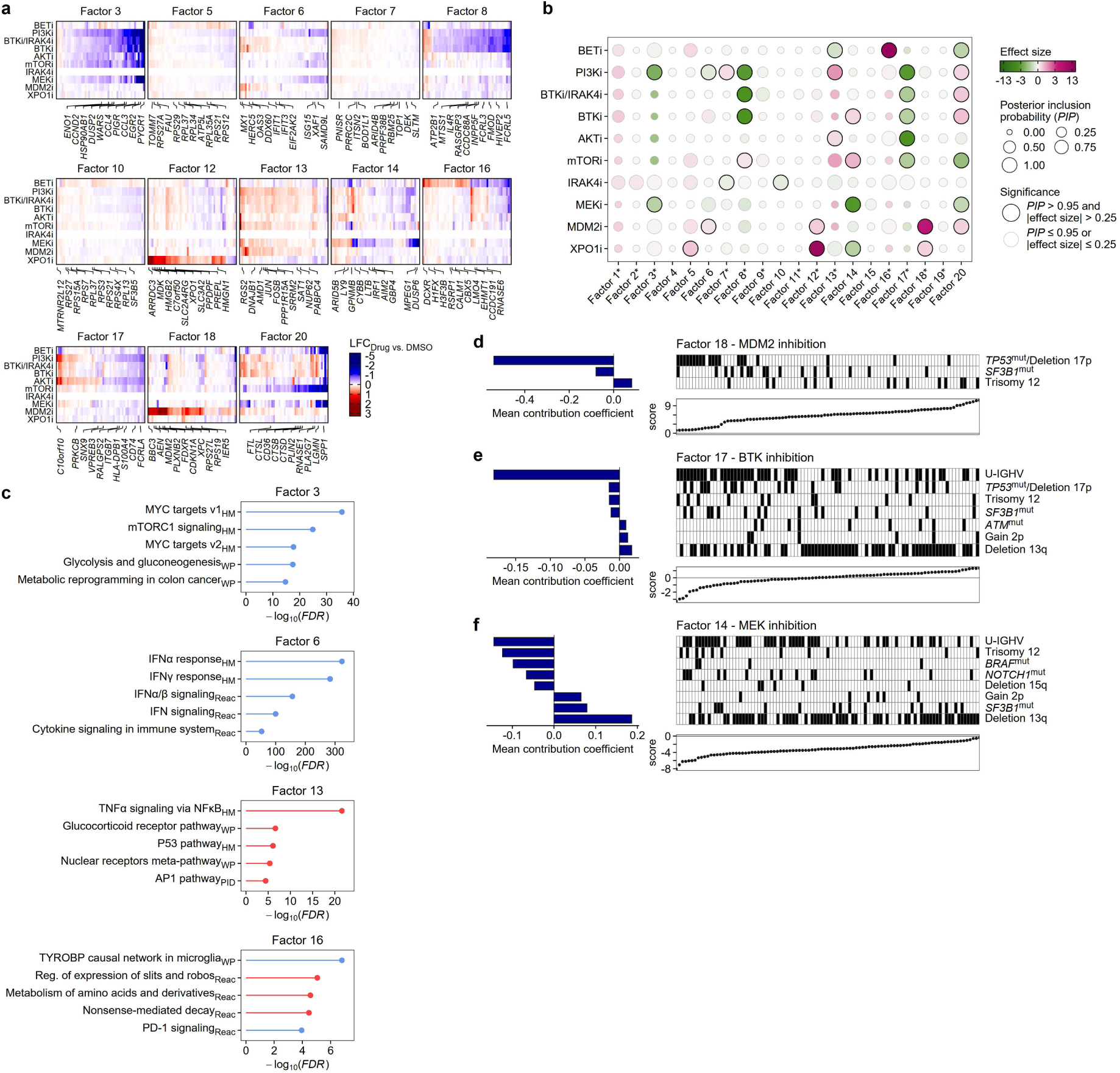
Guided sparse factor analysis of drug effects. **a**, Heatmap shows drug effects for 50 genes with the highest gene weights for factors with significant loadings. Top 10 genes with the highest gene weights for each factor are labelled. **b**, Estimated effects of drug perturbations on co-regulated gene modules represented by 20 latent factors. The posterior inclusion probability (*PIP*) denotes the probability that a perturbation affects the factor loading. Factors with significant loadings (*PIP* > 0.95, |effect size| > 0.25) are shown. Asterisks denote factors for which the sign was inverted to reflect the direction of the drug effects. **c**, GSEA with the gene weights for each factor using hallmark and canonical pathways gene sets. Gene sets were filtered by *FDR* (< 1%) and number of enriched genes (> 10). The sign of weights was reverted in accordance to (b). **d**, Influence of genetic disease drivers on the contribution of the MDM2 inhibitor effect in factor 18, **e**, BTK inhibitor effect in factor 17, and **f**, MEK inhibitor effect in factor 14 computed by multiple linear regression.

**Suppl. Fig. 7:**
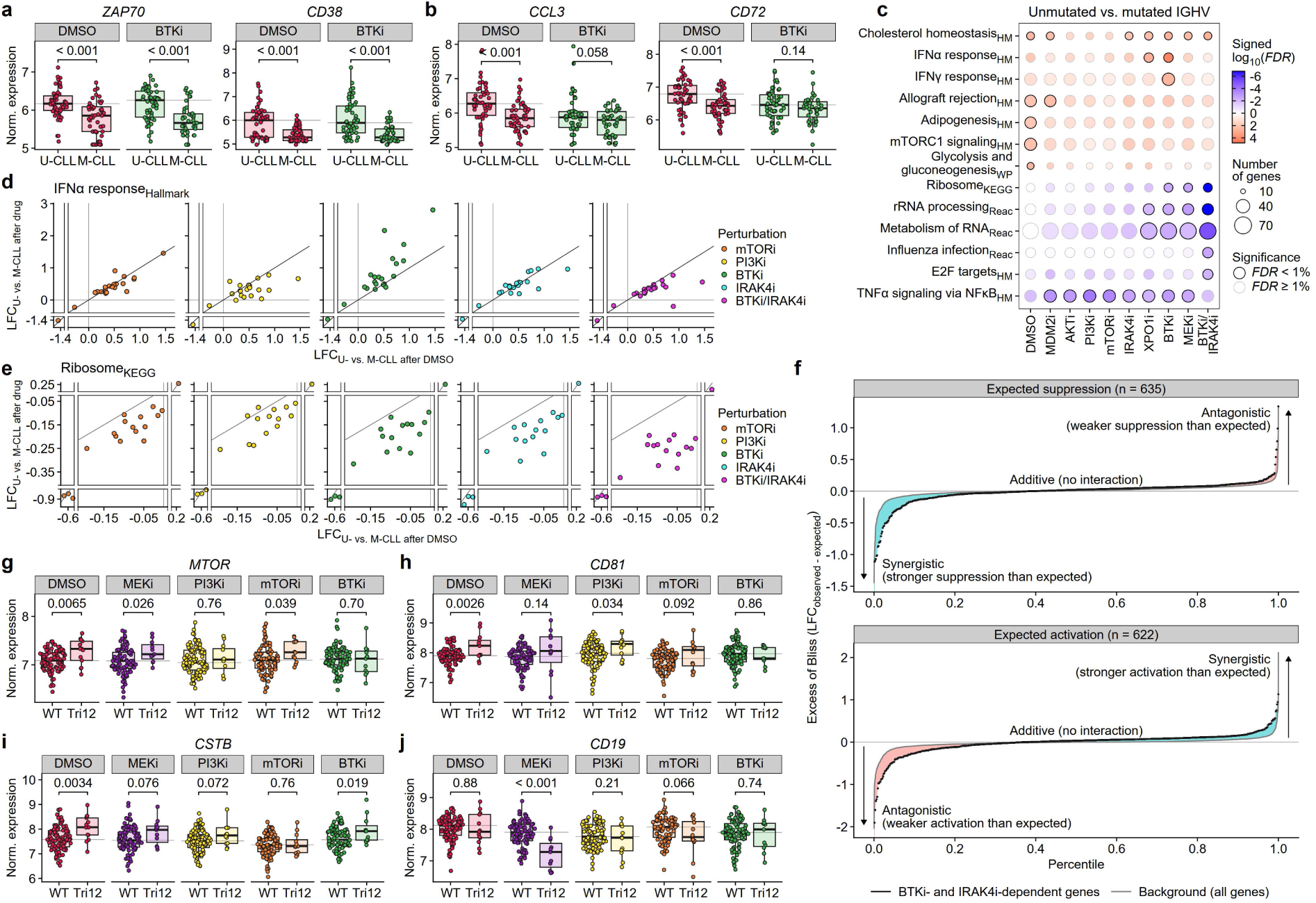
Pathway dependency of expression signatures and disease drivers. **a**, Normalised expression of *ZAP70 and CD38* by IGHV status in DMSO or BTK inhibitor-treated samples. **b**, Normalised expression of *CCL3* and *CD72* by IGHV status in DMSO or BTK inhibitor-treated samples. Significance levels were calculated using two-tailed Student’s *t*-test and shown as nominal *P*-values. **c**, GSEA with genes differentially expressed between U- and M-CLL (10% *FDR* in > 90% of bootstrapping performed to increase stringency) for each treatment using hallmark and canonical pathways gene sets. Gene sets were filtered by *FDR* (< 1%) and number of enriched genes (> 10). **d**, Differential expression of IGHV signature genes within the IFN response and **e**, ribosome gene set with BTKi, PI3Ki, mTORi, IRAK4i and BTKi/IRAK4i compared to DMSO. **f**, Expected independent additive effect from single BTK and IRAK4 inhibition against the observed effect from the BTK/IRAK4 inhibitor combination for down- and up-regulated BTK, IRAK4 and BTK/IRAK4 inhibitor-dependent genes (black dots). The expected independent effect was calculated using the Bliss independence model (LFC_expected_ = LFC_BTKi_ + LFC_IRAK4i_ - LFC_BTKi_ * LFC_IRAK4i_) and compared to the observed effect (LFC_BTKi/IRAK4i_). The gray dots depict the background distribution of expected additive effect for all genes (n_Down_ = 8,821, n_Up_ = 8,040). Shaded areas indicate increased synergistic (blue) or antagonistic (red) effects for BTK, IRAK4 and BTK/IRAK4 inhibitor-dependent genes compared to all genes. **g**, Normalised expression of dynamic BCR-dependent genes MTOR and **h**, CD81, and **i**, mTOR-dependent gene *CSTB* divided by trisomy 12 status in DMSO or drug-treated samples. **j**, Normalised expression of MEK-induced trisomy 12 signature gene *CD19* divided by trisomy 12 status in DMSO or MEK inhibitor-treated samples. Significance levels were calculated using two-tailed Student’s *t*-test and shown as nominal *P*-values.

**Suppl. Fig. 8:**
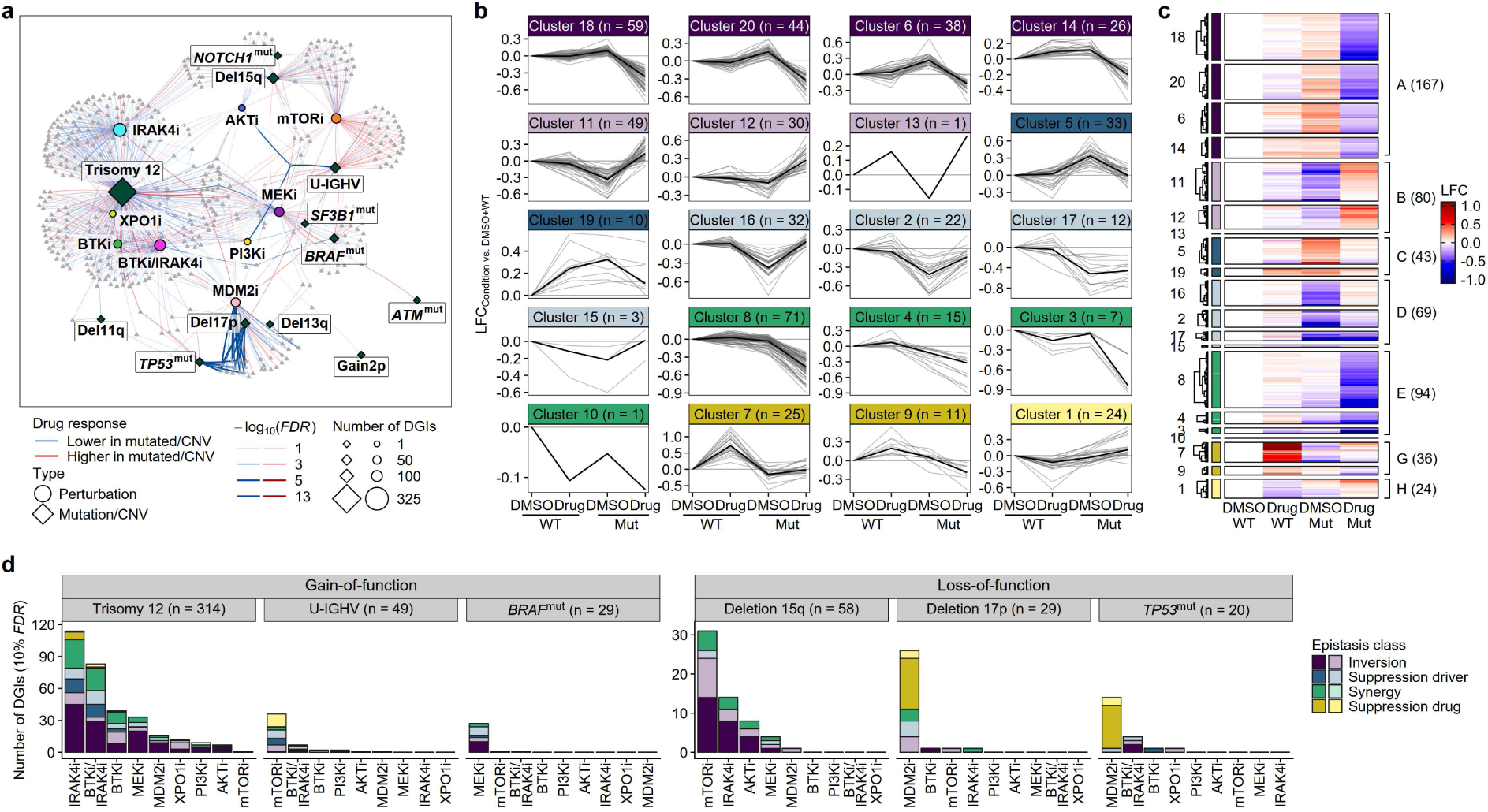
Interaction of perturbation effects and disease drivers. **a**, Pathway connectivity network of all DGI genes shown as a bipartite graph. Genes, drugs and disease drivers are connected if a differential drug response was observed in the presence of the mutation (*FDR* < 10%). **b**, Cluster assignment of DGIs based on gene expression in the presence of either drug, mutation/CNV or both compared to WT/DMSO samples by Fuzzy C-Means Clustering. The thick black line depicts the median LFC of all genes from the respective cluster. **c**, Heatmap of relative gene expression of DGIs in the presence of either drug, mutation/CNV or both compared to WT/DMSO samples by epistasis class. **d**, Number of target genes with significant DGIs for each perturbation, and all recurrent drivers with more than 5 cases by epistasis class (*FDR* < 10%).

**Suppl. Fig. 9:**
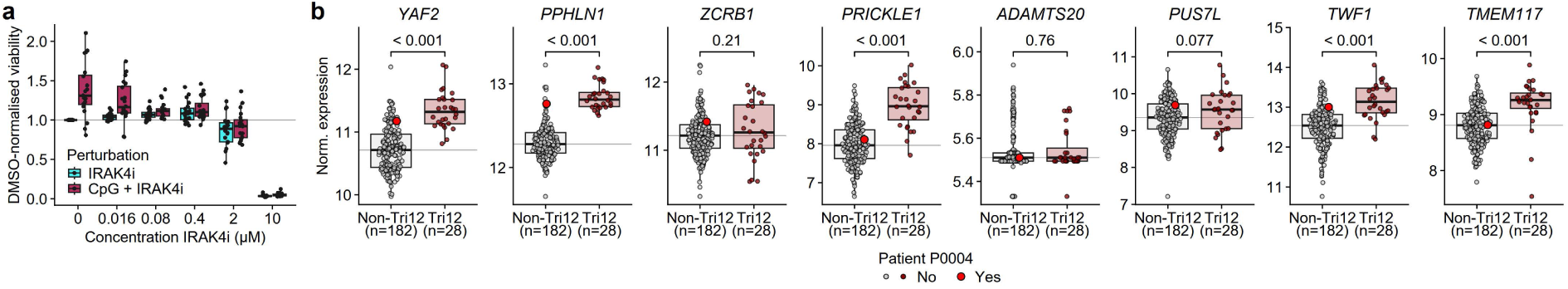
IRAK4 is a disease driver targeted by trisomy 12. **a**, Dose-response curves of 20 patients treated with IRAK4 inhibitor and IRAK4 inhibitor combined with 1 µg/mL CpG-ODN for TLR9 stimulation. **b**, Gene expression of genes on the amplified locus chr12:42,538,673-chr12: 44,229,999 (hg19) by trisomy 12 status for the CLL cohort (n = 210) from Lütge et al.^13^. Significance levels were calculated using two-tailed Student’s *t*-test and shown as nominal *P*-values.

## Supplementary methods figures

**Suppl. Methods Fig. 1:**
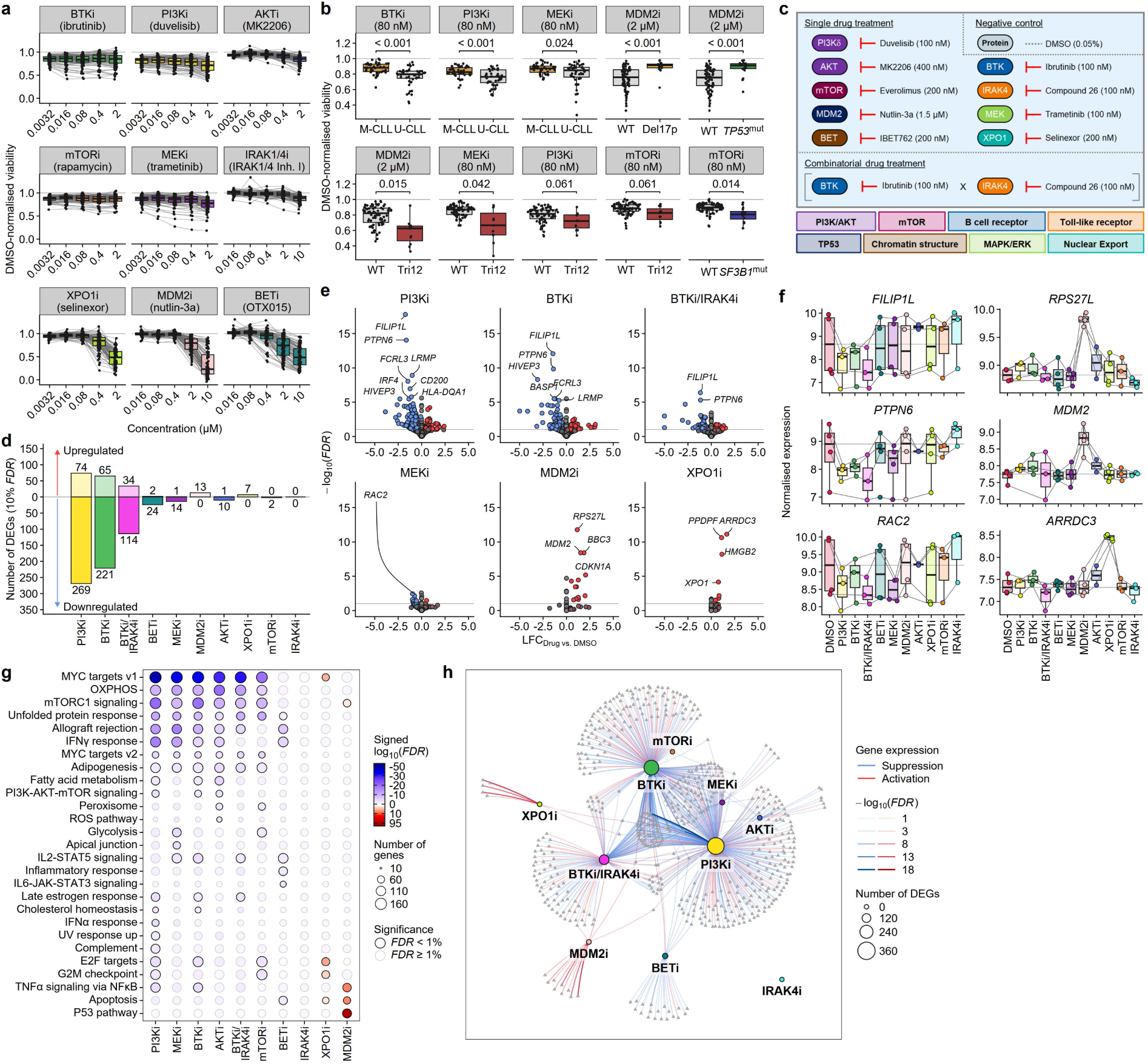
Design and pilot drug screening using shallow QuantSeq. **a**, Dose-response curves of drug response in CLL patients (n = 88) (viability drug screen from Lu et al.^12^). **b**, Significant genetic association with drug response in CLL samples (*FDR* < 10%) for aberration with more than 5 cases. Significance levels were calculated using two-tailed Student’s *t*-test and adjusted for multiple testing. **c**, Overview of treatments and concentrations. **d**, Number of DEGs between 4 drug- and DMSO-treated CLL samples. **e**, Volcano plot summarising the differential gene expression after PI3K, BTK, combined BTK/IRAK4, MEK, MDM2 and XPO1 inhibitor treatment. **f**, Gene expression of exemplary genes with treatment-specific down- and upregulation. **g**, GSEA with DEGs after each treatment (all genes) using hallmark gene sets. Gene sets were filtered by *FDR* (< 1%) and number of enriched genes (> 10). **h**, Pathway connectivity network shown as a bipartite graph. Genes and drugs are connected if the drug perturbation leads to differential expression (10% *FDR*).

**Suppl. Methods Fig. 2.**
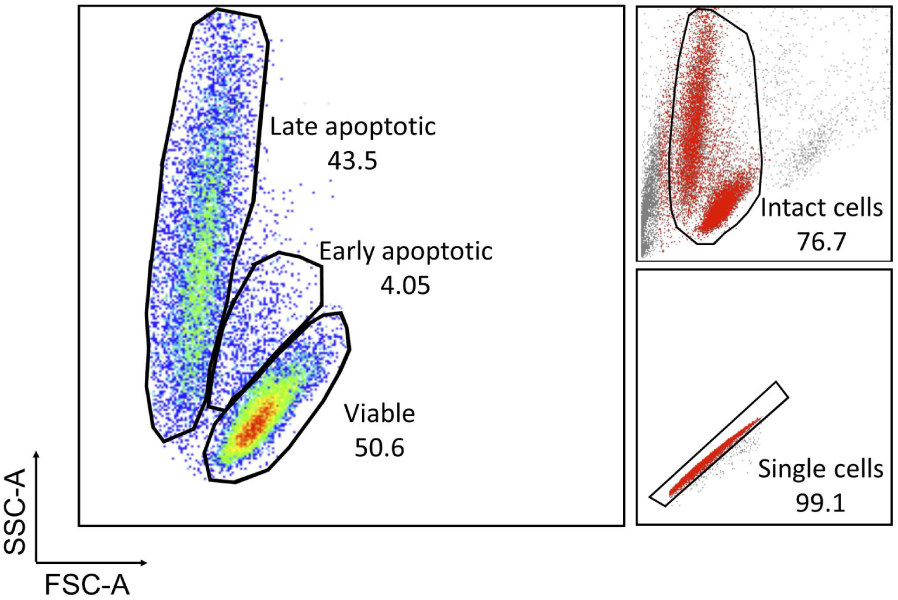
Gating strategy for flow cytometric assessment of cell viability. Cells were pre-gated by excluding debris and doublet cells. Viable and dead cells were then gated based on cell size (FSC-A) and granularity (SSC-A).

**Suppl. Methods Fig. 3.**
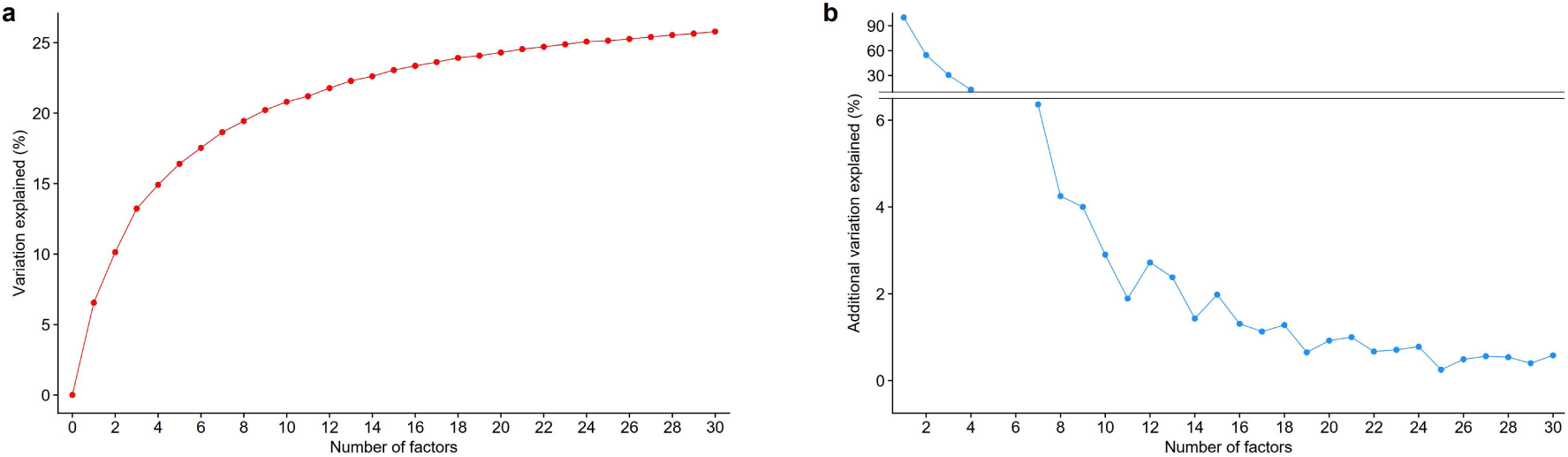
Guided sparse factor analysis model evaluation. **a**, Proportion of variation explained by each GSFA model with 1 to 30 factors. **b**, Percentage of additional variation explained by each GSFA model compared to the previous model with the highest explained variation. The model with 20 factors was selected for further analysis.

